# Complex Formation of Immunoglobulin Superfamily Molecules Side-IV and Beat-IIb Regulates Synaptic Specificity in the *Drosophila* Visual System

**DOI:** 10.1101/2023.03.27.534487

**Authors:** Jiro Osaka, Arisa Ishii, Xu Wang, Riku Iwanaga, Hinata Kawamura, Atsushi Sugie, Satoko Hakeda-Suzuki, Takashi Suzuki

## Abstract

Neurons establish specific synapses based on the adhesive properties of cell-surface proteins, while also retaining the ability to form synapses in a relatively non-selective manner. However, the comprehensive understanding of the underlying mechanism reconciling these opposing characteristics remains incomplete. Here, we have identified Side-IV/Beat-IIb, members of the *Drosophila* immunoglobulin superfamily, as a novel combination of cell-surface recognition molecules inducing synapse formation. The Side-IV/Beat-IIb combination transduces bifurcated signaling with Side-IV’s co-receptor, Kirre, and a synaptic scaffold protein, Dsyd-1. Genetic experiments and subcellular protein localization analyses showed the newly characterized Side-IV/Beat-IIb/Kirre/Dsyd-1 complex to have two essential functions. First, it narrows neuronal binding specificity through Side-IV/Beat-IIb extracellular interactions. Second, it recruits synapse formation factors, Kirre and Dsyd-1, to restrict synaptic loci and inhibit miswiring. This dual function explains how the combinations of cell-surface molecules enable the ranking of preferred interactions among neuronal pairs to achieve synaptic specificity in complex circuits *in vivo*.

## INTRODUCTION

Neurons possess an intriguing capability called synaptic specificity that enables them to selectively establish synapses with specific neurons while excluding others.^1, 2^ During development, neurons stochastically extend filopodia, searching for synaptic partners.^3–5^ Synapse formation begins only when filopodia find a synaptic partner through specific molecular interactions.^2, 3^ Through the study in cultured cells, several cell-surface proteins have been found to have mutual binding abilities that can induce synapse formation.^6^ However, *in vivo* investigations have revealed that neurons possess the ability to form a consistent number of synapses indiscriminately, while simultaneously exhibiting a hierarchical preference for synaptic connections.^7, 8^ The comprehensive molecular mechanisms underlying the interplay between synaptic specificity and preferences in neurons remain largely unexplored.

The *Drosophila* visual system consists of a compound eye and four optic ganglia, compactly composed of approximately 200 diverse neurons with well-documented connections.^9, 10^ In addition, single-cell analyses in the last few years have revealed cell-type and time-specific gene expression profiles of the *Drosophila* visual system, which have been compiled into a database such as Gene Expression Omnibus.^11–15^ This makes the *Drosophila* visual system an ideal model for connectome and transcriptome-based analyses. Several transmembrane protein pairs with binding interactions have been reported to be expressed between synaptic partners.^13, 16–18^ The most studied is the DIP-Dpr families of the immunoglobulin superfamily (IgSF), which form a heterophilic or homophilic interaction network.^16, 19^ In recent years, it has been shown that DIP-Dpr protein interactions are involved in neural connections, axon guidance, and cell death signaling.^7, 17, 18, 20–24^ Another member of IgSF, the Beat-Side families have similar properties to DIP-Dpr, although their functions are less well understood. Therefore, we considered the Beat-Side families as candidate molecules potentially involved in neural circuit formation.

Both *beaten path Ia* (*beat-Ia*) and *sidestep* (*side*) were identified by the genetic screening of mutants showing abnormal axon guidance in neuromuscular junctions (NMJ).^25–27^ The *beat-Ia* and *side* mutants had similar phenotypes and were shown to interact genetically and biochemically.^16, 28, 29^ Genomic and interactome analyses revealed that *beat-Ia* and *side* have 14 and 8 paralogs, respectively, with a heterophilic interaction network.^16, 29–32^ Furthermore, single-cell RNAseq data showed that Beat-Side families are expressed in a cell-specific manner throughout the nervous system.^11, 13^ Although several works have identified the functions of Beat-Ia-Side interactions to be axonal guidance of motor axons, motor function, and leg nerve development,^28, 33–35^ the functions of other Beat-Side families are yet to be elucidated, especially in the central nervous system.

In the present study, we report a novel cell-surface protein combination for synaptic specificity between Side-IV and Beat-IIb. Overexpression of *side-IV* can induce synapse formation in the R7 photoreceptors of *Drosophila*. Triggered by *trans*-interactions with Beat-IIb, Side-IV creates a cluster complex that could transduce bifurcated signaling to induce synapses with its co-receptor, Kirre, and a synaptic scaffold protein, Dsyd-1. Contrary to our assumption, our *side-IV* loss-of-function analysis revealed more ectopic synapse formation in the distal parts of lamina monopolar neurons. Combining the analyses of the loss- and gain-of-function from one side and the subcellular protein localization from another side, we were able to demonstrate that a complex formed of Side-IV, Beat-IIb, Kirre, and Dsyd-1 not only narrows neuronal binding specificity but also recruits synapse formation factors Kirre and Dsyd-1 to restrict synapse formation loci and inhibit miswiring. Consequently, we proposed a mechanism whereby the specific combinations of cell-surface molecules decide the hierarchy of preference among neuronal pairs in a complex circuit *in vivo*.

## RESULTS

### Side-IV and Beat-IIb are a New Cell-Surface Combination for Synaptic Induction

To identify the cell-surface protein(s) able to induce synapse formation, we focused on the Beat-Side families and their functional roles. Accordingly, we analyzed the R7 photoreceptor, which extends its axons to the medulla and predominantly forms synapses with the dendrites of Dm8 in the M4–M6 layer.^4,10, 36, 37^ To visualize the presynaptic structures of R7 in a cell-specific manner, we applied the synaptic tagging with recombination (STaR) technique, which allows the active zone protein, Bruchpilot (Brp), to be labeled using the R7-specific FLPase, 20C11-FLP (**Figure 1A**).

**Figure 1.**
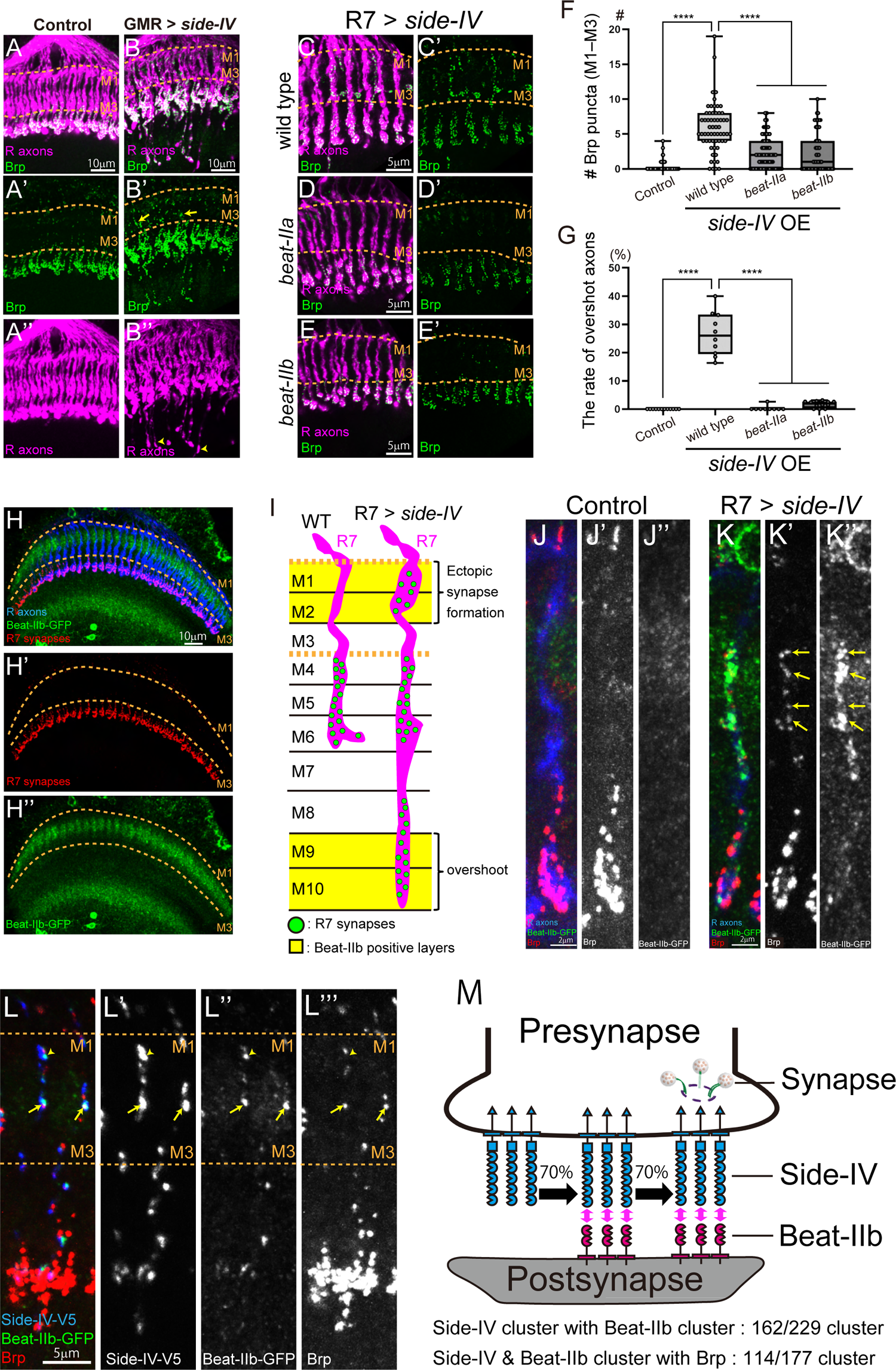
Interaction of Side-IV and Beat-IIb is Instructive for Synaptic Induction. (A–B’’) Ectopic synapse formation (arrow) and overshot axons (arrowheads) of R7 in GMR > *side-IV* overexpression flies (B–B’’). Controls are shown in (A–A’’). Synapses are visualized by STaR technique^37^ combining with R7-specific-FLPase (20C11-FLP). Photoreceptor axons (R axons) are stained by mAb24B10. Dotted lines indicate M1–M3 layer. (C–E’) Ectopic synapse formation of R7 in *side-IV* overexpression (OE) by R7-specific-gal4 (GMR-FsF-gal4 + 20C11-FLP) in various mutant backgrounds: wild type (C and C’), *beat-IIa* (D and D’), and *beat-IIb* mutants (E and E’). Miswiring phenotypes were suppressed significantly in *beat-IIa* or *beat-IIb* mutants. (F) Quantification of the number of the ectopic synapses (n > 60 axons). (G) Quantification of the rate of overshot axons (n >10 brains). (H–H’’) Axonal and synapse morphology of R7 and Beat-IIb-MiMIC-GFP localization in the medulla. (I) Phenotypic scheme of *side-IV* overexpression. Miswiring phenotypes occurred in Beat-IIb positive layers. (J–K’’) Punctate signal of Beat-IIb-MiMIC-GFP in R7 > *side-IV* overexpression. Control (J–J’’), *side-IV* overexpression (K–K’’). Arrows indicate punctate Beat-IIb signal where Brp were induced adjacently. (L–L’’’) Mutual localization of Side-IV-V5, Beat-IIb-MiMIC-GFP, and Brp of R7. Side-IV-V5 is expressed specifically in R7. Side-IV/Beat-IIb cluster with (arrows) and without (arrowheads) Brp. (M) Summary scheme of Side-IV clusters, Beat-IIb clusters, and synaptic induction. Statistical treatment of the number synapses and the rate of overshot axons were analyzed with the Mann-Whitney test. In this and all following statistical significance. n.s. *P* > 0.05, * *P* ≤ 0.05, ** *P* ≤ 0.01, *** *P* ≤ 0.001, **** *P* ≤ 0.0001. See also **Figure S1**.

Gene knockdown (**Figures S1A** and **S1B**) and overexpression (**Figures S1C** and **S1D**) experiments with Beat-Side family members in R7 showed that overexpression of *side-IV* results in the induction of ectopic synapses in the M1–M3 layer of the medulla (**Figures 1B** and **1B’**, yellow arrows, and **S1C**), as well as overshooting of R7 axons toward the M10 layer (**Figures 1B** and **1B’’**, yellow arrowheads). These phenotypes indicated that R7 was miswired in regions other than M4–M6, its usual target layer, due to the expression of *side-IV*. It was noteworthy that the endogenous tags of Side-IV that revealed R7 did not endogenously express *side-IV* (**Figures S1E–I’**). Given the biochemical interaction identified between Side-IV and Beat-IIa/b,^16, 29^ we next investigated the contribution of Side-IV-Beat-IIa/b interactions to the observed miswiring phenotypes. Both synaptic induction and overshoot phenotypes generated by *side-IV* overexpression were significantly suppressed in a *beat-IIa* or *beat-IIb* mutant background (**Figures 1C–G**). We used the endogenous Beat-IIb-GFP marker^38^ to determine the expression pattern of the beat ligands and found that *beat-IIb* was specifically expressed in the M1–M2 and M9–M10 layers, where the miswiring phenotypes had been detected (**Figures 1H–I**). The expression of *side-IV* caused the Beat-IIb-GFP signal to become punctate (**Figures 1J–K’’**) and presynaptic Brp puncta was induced adjacently (**Figures 1K’** and **1K’’**, yellow arrows). This punctate signal was indicative of the clustering of transmembrane proteins, a common phenomenon observed with other synapse-organizing molecules.^39^ The concurrent visualization of Side-IV, Beat-IIb, and Brp also revealed clustered signals of Side-IV, approximately 70% of which co-localized with Beat-IIb clusters (**Figures 1L–L’’’**). Furthermore, approximately 70% of Side-IV/Beat-IIb clustered signals co-localized with ectopic synapses (**Figures 1L–L’’’**, yellow arrows). These findings strongly suggest that Side-IV induced the clustering of Beat-IIb, which, in turn, could drive the induction of synapse formation (**Figure 1M**). Therefore, we identified Side-IV/Beat-IIb as a cell-surface protein combination potentially capable of inducing synapses by establishing cluster complexes.

### Side-IV Mediates Cluster Complex Formation via Extracellular Domain and Transduces Signals by Co-Receptors and Cytoplasmic Domain

To uncover the synaptic induction mechanism of Side-IV, we examined the contributions of its individual domains. Side-IV comprises five extracellular Ig domains (Ig1–5), a fibronectin type III domain (FnIII), and a cytoplasmic domain (Cyto) with a PDZ-binding (PDZb) motif (**Figures 2A** and **S2A**). By generating domain-deficient variants and chimeras of Side-IV with a nonfunctional membrane protein, mCD8 (**Figures 2A** and **2B**), we assessed the Side-IV/Beat-IIb clustering and ectopic synapse/overshoot induction phenotypes in R7 (**Figures 2C–K** and **S2B–K**). The deletion of the extracellular Ig domain abolished the ability of Side-IV to induce the phenotypes (**Figures S2B–E** and **S2I–K**), confirming that an interaction between Side-IV and Beat-IIb occurs in *trans*, as observed in *Drosophila* S2 cell cultures (**Figures S2L–Q**).

**Figure 2.**
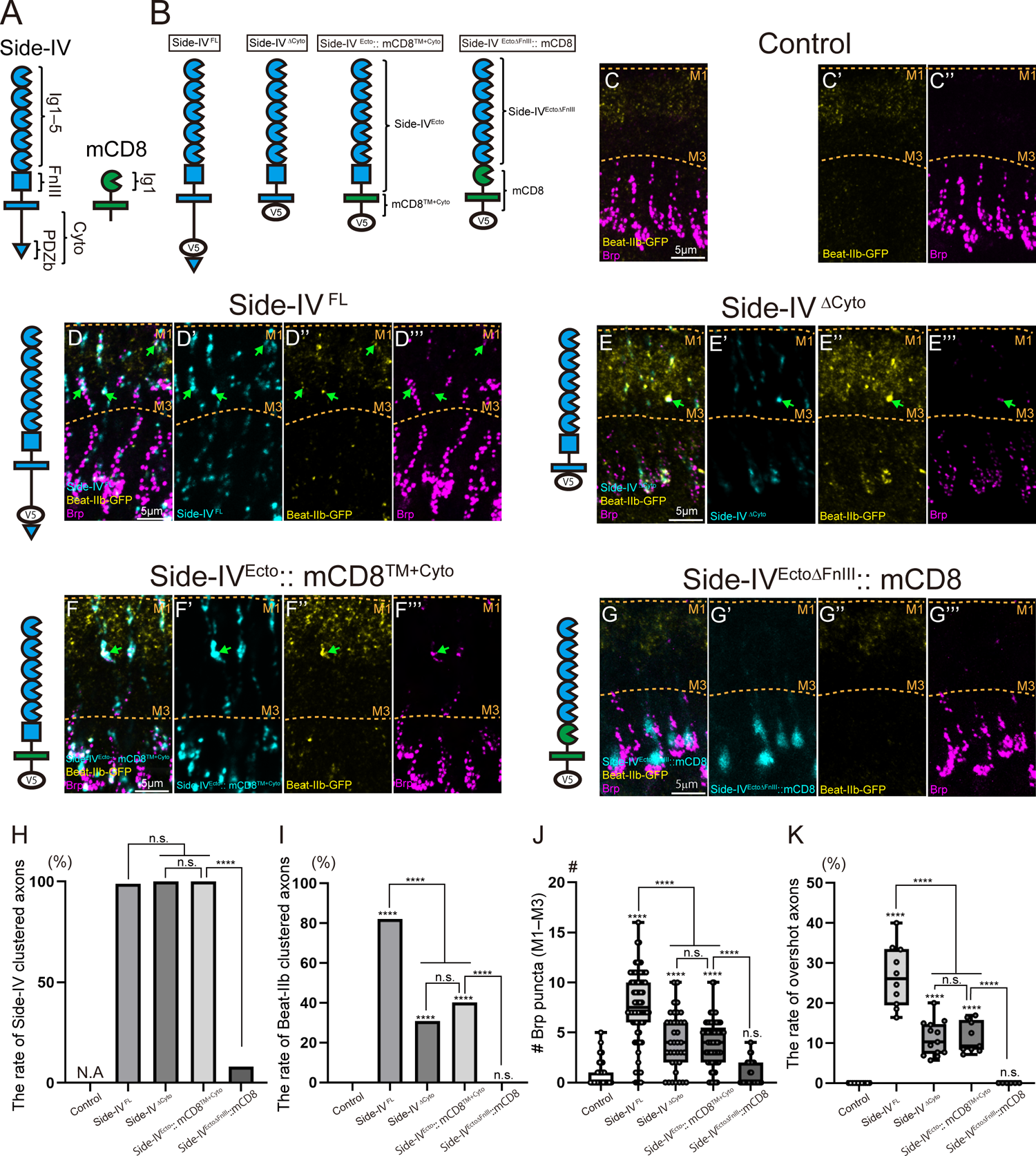
Cytoplasmic and FnIII Domains of Side-IV Contribute for Ectopic Synapse Formation. (A) Domain structure of Side-IV and mouse CD8 (mCD8). (B) Variants of Side-IV chimera used in this Figure. Side-IV ^FL^ has a V5 sequence before the PDZ binding (PDZb) motif. Other variants have a V5 sequence on their C-terminals. (C–G’’’) Side-IV/Beat-IIb cluster and ectopic synapse formation on each Side-IV variants. Arrows indicate Side-IV/Beat-IIb clusters that induced ectopic synapse. Variants of *side-IV* are overexpressed by R7-sepcific-gal4 (GMR-FsF-gal4 + 20C11-FLP) and stained by α-V5. Brp are visualized by STaR technique with Brp-FsF-mCherry and 20C11-FLP. Merge (C, D, E, F, G), Side-IV-variants-V5 (cyan, D’, E’, F’, G’), Beat-IIb-MiMIC-GFP (yellow, C’, D’’, E’’, F’’, G’’), Brp-mCherry (magenta, C’’, D’’’, E’’’, F’’’, G’’’). Mis-localization Side-IV ^EctoΔFnIII^::mCD8 indicates FnIII domain is necessary for Side-IV clustering. Dotted lines indicate M1–M3 layer. (H) Quantification of the rate of Side-IV-variants-V5 clustered axons (n > 28 axons). (I) Quantification of the rate of Beat-IIb-MiMIC-GFP clustered axons (n > 47 axons). (J) Quantification of the number of the ectopic synapses (n > 32 axons). (K) Quantification of the rate of overshot axons (n > 6 brains). Statistical treatment of the rate of Side-IV-variants-V5 or Beat-IIb-MiMIC-GFP clustered axons were analyzed with the Chi-square test. Statistical treatment of the number of the ectopic synapses and the rate of overshot axons were analyzed with the Mann-Whitney test. See also Figures S2 and S3.

The overexpression of the deletion construct of the cytoplasmic domain (Side-IV^ΔCyto^) in R7 resulted in a milder, yet still robust phenotype in both synapse induction and overshoot axons (**Figures 2E–E’’’**, **2J**, and **2K**). This robustness in synapse-inducing ability suggests that cytoplasmic domain may not be essential, but rather, specific enhancement in cell affinity could induce synapses. To explore whether just the enhancement of physical affinities between neurons could induce synapse formation in this system, we introduced homophilic interactions of epithelial cadherins or human-derived sidekick cell adhesion molecule 2 (SDK2) between R7 and postsynaptic neurons (**Figures S3A–D**). However, the results were negative. Notably, the applied gal4 drivers^40^ were sufficient to induce synapses when the known synapse-inducing proteins, Nrxs-Nlgs, were expressed (**Figures S3E–J**). This indicated that increasing the affinity was insufficient to induce synapse formation. Therefore, we inferred that the Side-IV/Beat-IIb cluster complex may transduce signals via its co-receptors. Additionally, signaling through the PDZb motif contributed to synaptic induction, as evidenced by weaker synapse induction with Side-IV^ΔPDZb^ compared to full-length Side-IV (**Figures S2C** and **S2H–I**).

The critical role of the FnIII domain became apparent when we compared the two chimeric proteins, Side-IV^Ecto^::mCD8^TM+Cyto^ and Side-IV ^EctoΔFnIII^::mCD8. The key difference between these two constructs is the presence or absence of the FnIII or mCD8’s Ig domain (**Figure 2B**). Like Side-IV^ΔCyto^, Side-IV^Ecto^::mCD8^TM+Cyto^ (with FnIII) exhibited robust induction of Side-IV clusters and synapses in R7 (**Figures 2E–F’’’** and **2H–K**). In contrast, Side-IV^EctoΔFnIII^::mCD8 (without FnIII) showed no cluster signal in the M1–M3 layer of R7 and failed to induce synapses (**Figures 2G–K**). Importantly, Side-IV^EctoΔFnIII^::mCD8 maintained binding affinity to Beat-IIb, since *Side-IV^EctoΔFnIII^::mCD8* transfected cultured cells aggregated strongly with *beat-IIb* transfected cells, comparable to *side-IV ^FL^* (**Figures S2P–Q**). These findings indicate that the FnIII domain is indispensable for Side-IV clustering and, consequently, for synapse induction.

Taken together, we show that the extracellular Ig and FnIII domains of Side-IV are essential for establishing the Side-IV/Beat-IIb cluster complex, and the cytoplasmic domain of Side-IV and/or its co-receptors transduce signals for synapse organization.

### Kirre Can Form a Complex with Side-IV via the Extracellular Domain as a Co-Receptor

To identify the co-receptor of Side-IV, we focused on transmembrane proteins that could enhance Side-IV signaling. Among these candidates, the immunoglobulin superfamily protein Kirre^41^ exhibited the ability to enhance the phenotypes of Side-IV (**Figures 3A–F**). Kirre expression resulted in a doubling of the Side-IV overshoot phenotype but Kirre alone did not induce the overshoot (**Figure 3F**), indicating a synergistic rather than additive effect. To examine the impact of the cytoplasmic domains on this synergistic effect, we analyzed the overshoot phenotype and found that, without their cytoplasmic domains, the synergy between Side-IV and Kirre was minimal (**Figures 3G–P**). However, a strong synergy was observed when the cytoplasmic domain of either was present (**Figures 3L**, **3N**, and **3P**), suggesting that Kirre acts as a co-receptor, enhancing Side-IV signaling through its cytoplasmic domain.

**Figure 3.**
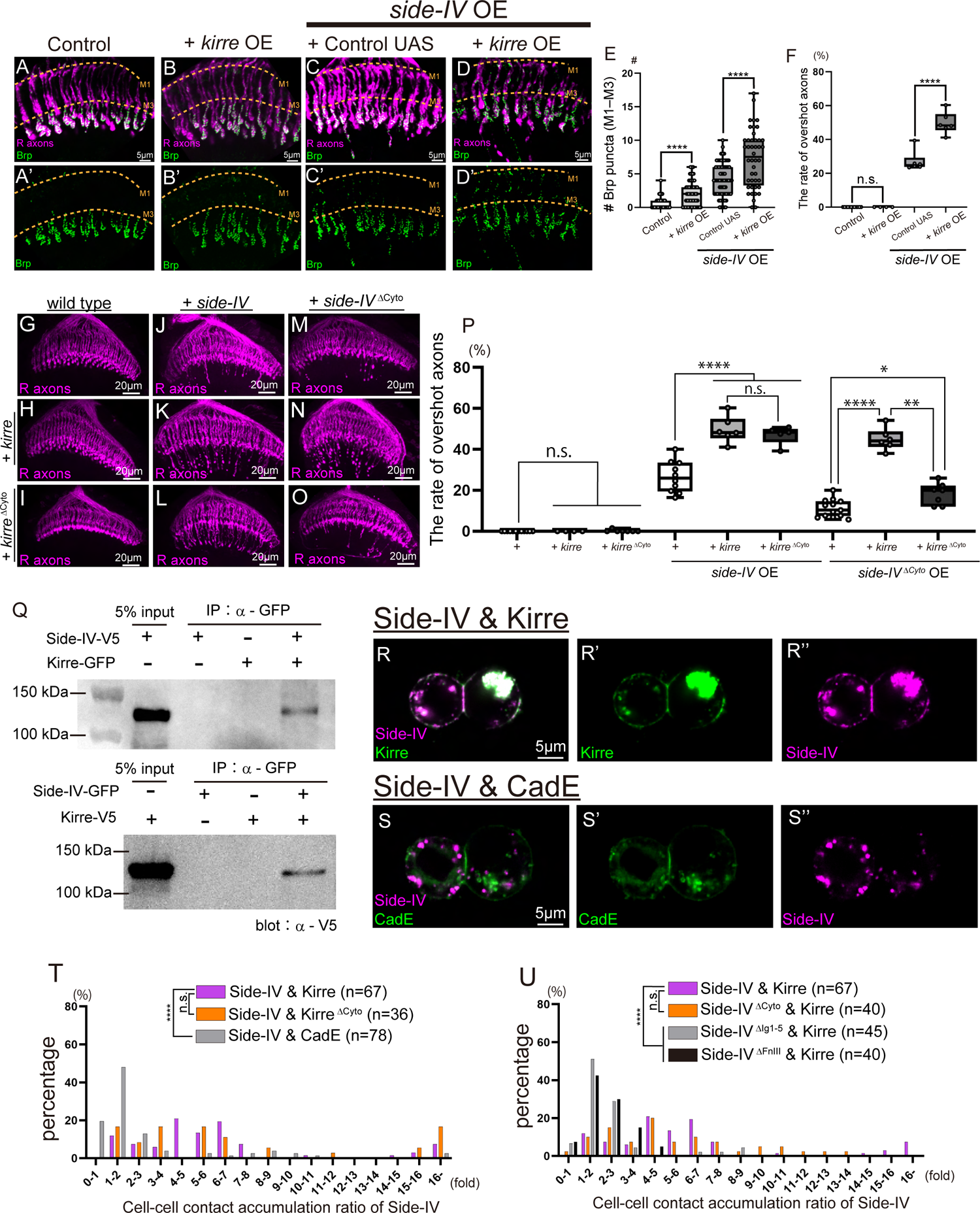
Kirre Interacts with Side-IV through the Extracellular Domain and Works as a Co-receptor. (A–D’) Ectopic synapse formation and overshoot phenotypes of R7 in GMR > *side-IV* and/or *kirre* overexpression (OE) flies. Control (A and A’), *kirre* OE (B and B’), Control UAS (*luciferase* RNAi) + *side-IV* OE (C and C’), *kirre* + *side-IV* OE (D and D’). R7 synapses are visualized by STaR technique. R axons are stained by mAb24B10. Dotted lines indicate M1–M3 layer. (E) Quantification of the number of the ectopic synapses (n > 44 axons). (F) Quantification of the rate of overshot axons (n > 4 brains). (G–O) Photoreceptors in medulla with co-expression of *side-IV* and *kirre* variants. Control (G), *kirre* OE (H), *kirre ^ΔCyto^*OE (I), *side-IV* OE (J), *side-IV* + *kirre* OE (K), *side-IV*+*kirre ^ΔCyto^*OE (L), *side-IV ^ΔCyto^*OE (M), *side-IV ^ΔCyto^*+ *kirre* OE (N), *side-IV ^ΔCyto^*+ *kirre ^ΔCyto^*OE (O). (P) Quantification of the rate of overshot axons (n > 4 brains). (Q) Result of co-immunoprecipitation for Side-IV and Kirre. S2 cell are transfected with Actin5C-gal4 and UAS-*side-IV*-V5/GFP and/or UAS-*kirre*-GFP/V5. Immunoprecipitation (IP) and immunoblotting (blot) were performed with α-GFP and α-V5, respectively. The bands between 100 kDa and 150 kDA were detected only when cells expressed both *side-IV* and *kirre*. Molecular weights of Side-IV-V5 and Kirre-V5 predicted by Bioinfomatics.org are 111.58 kDa and 102.53 kDa, respectively. (R–S’’) Protein accumulation at cell boundaries in transiently transfected S2 cells subjected to aggregation assay. S2 cells are co-transfected Actin5C-gal4 with UAS-*side-IV*-mCherry and UAS-*kirre*-GFP (R–R’’), UAS-*side-IV*-mCherry and UAS-*cadE*-GFP (S–S’’). Side-IV showed co-accumulation with Kirre accumulated cell contact sites. (T and U) Distribution of accumulation ratio of Side-IV co-transfected with Kirre, Kirre ^ΔCyto^, or CadE (T), Side-IV variants co-transfected with Kirre (U). Sample size is indicated (n). Cell-cell contact accumulation ratio is the ratio of the average fluorescence intensity of cell-cell contact sites to the average fluorescence intensity of cell membrane which does not contact. Statistical treatment of the number of synapses and the rate of overshot axons were analyzed with the Mann-Whitney test. Distribution of the cell-cell contact accumulation ration was analyzed with the unpaired t-test. See also **Figure S4**.

To demonstrate the biochemical interaction between Side-IV and Kirre, we successfully co-immunoprecipitated Side-IV with Kirre, and vice versa, using co-transfected cultured cells (**Figure 3Q**). To confirm that the interaction occurred in *cis*, we further employed the cell culture system. Initially, we showed that Side-IV and Kirre did not induce cell aggregation via *trans*-interactions in the cultured cells (**Figure S4**). Subsequently, we assessed their *cis*-interaction using cell-cell contact sites.^42, 43^ Side-IV co-accumulated with Kirre at the cell boundaries, where homophilic interaction of Kirre accumulated from both sides of the cell membranes (**Figures 3R–R’’**). Importantly, this co-accumulation was specific to Kirre, as the Side-IV accumulation did not occur with the accumulation of CadE at the cell-cell contact sites (**Figures 3S–S’’**). Furthermore, the co-accumulation required Side-IV’s extracellular domain but not its cytoplasmic domain (**Figures 3T** and **3U**). Collectively, these data suggest that Kirre functions as a co-receptor of Side-IV and, together, they form a Side-IV/Kirre complex established through *cis*-interaction and mediated by their extracellular domains.

### Dsyd-1 & Liprin-α From Complex with Side-IV and Act Downstream

The reduced synapse formation of Side-IV^ΔCyto^ compared to the full-length suggests the functional requirement of cytoplasmic signaling. To identify the downstream targets of Side-IV, we searched for the synapse-related molecules necessary for Side-IV-induced synapse formation. We found that RNAi knockdown of *dsyd-1* or *liprin-α* significantly suppresses synaptic induction by *side-IV* overexpression in R7 (**Figures 4A–F**), indicating that Dsyd-1 and Liprin-α function downstream of Side-IV.

**Figure 4.**
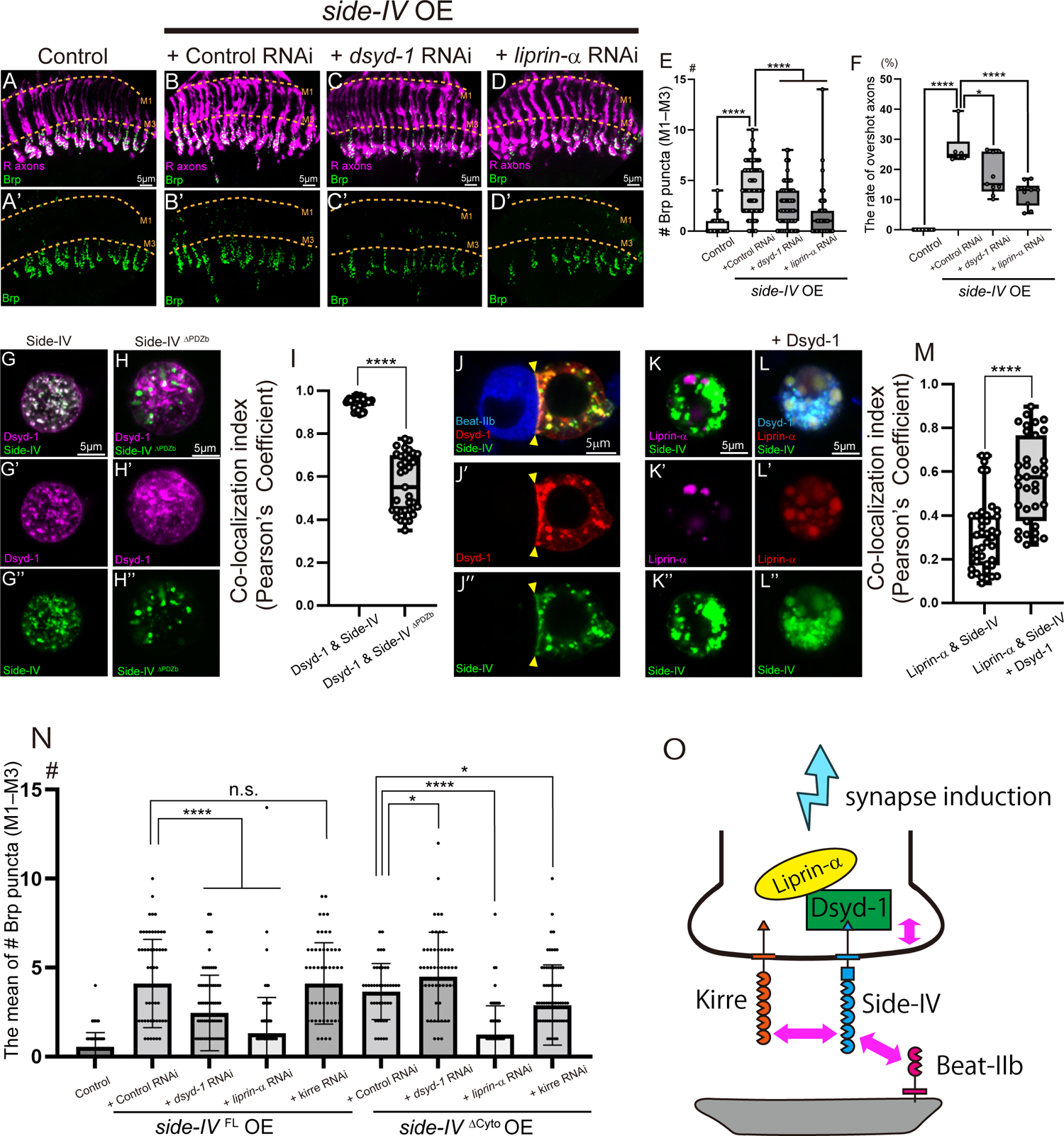
Side-IV Transduces a Signal with Dsyd-1 and Liprin-α Downstream. (A–D’) Ectopic synapse formation and overshoot phenotypes of R7 in GMR > *side-IV* with *dsyd-1* or *liprin-a* RNAi. Control (A and A’, same images as **3A** and **3A’**), Control RNAi (*luciferase* RNAi) + *side-IV* OE (B and B’, same images as **3C** and **3C’**), *dsyd-1* RNAi + *side-IV* OE (C and C’), *liprin-α* RNAi + *side-IV* OE (D and D’). R7 synapses are visualized by STaR method. R axons are stained by mAb24B10. Dotted lines indicate M1–M3 layer. (E) Quantification of the number of the ectopic synapses (n > 69 axons). (F) Quantification of the rate of overshot axons (n > 6 brains). (G–H’’) Localization of Side-IV and Dsyd-1. S2 cells were co-transfected with Actin5C-gal4, UAS-*dsyd-1*-mCherry, and UAS-*side-IV*-GFP or UAS-*side-IV ^ΔPDZb^*-GFP. Side-IV-GFP and Dsyd-1-mCherry (G–G’’), Side-IV ^ΔPDZb^-GFP and Dsyd-1-mCherry (H–H’’). Side-IV strongly co-localized with Dsyd-1 when PDZ binding motif exists. (I) Quantification of the co-localization (n > 35 images). (J–J’’) Localization of S2 cell co-transfected with Actin5C-gal4, UAS-*side-IV*-GFP (green), and UAS-*dsyd-1*-mCherry (red) aggregated with S2 cell co-transfected Actin5C-gal4 and UAS-*beat-IIb*-V5 (blue). Arrowheads represent the Side-IV accumulated cell contact sites where Dsyd-1 showed co-accumulation. (K–L’’) Localization of Side-IV and Liprin-α. S2 cells were co-transfected Actin5C-gal4 with UAS-*side-IV*-GFP, UAS-*liprin-a*-mCherry without (K–K’’) and with (L–L’’) UAS-*dsyd-1*-V5. Side-IV co-localized with Liprin-α when *dysd-1* is co-expressed. (M) Quantification of the co-localization (n > 35 images). (N) Quantification of the number of ectopic synapses (n > 43 axons) represented as mean ± SD. (O) Working model of synaptic induction by Side-IV. Statistical treatment of the number of synapses and the rate of overshot axons were analyzed with the Mann-Whitney test. Statistical treatment of the Co-localization index was analyzed with the unpaired t-test. See also **Figure S5**.

Using S2 cells, we examined the co-localization of Side-IV with Dsyd-1 and Liprin-α (**Figures 4G– M**). Side-IV exhibited substantial co-localization with Dsyd-1 (**Figures 4G–G’’** and **4I**). Since Dsyd-1 possesses a PDZ domain that interacts with synapse-organizing molecules such as Nrx-1,^44^ we hypothesized that Side-IV interacts with Dsyd-1 through Side-IV’s PDZb motif. The co-localization index between Side-IV and Dsyd-1 decreased by almost half in the absence of Side-IV’s PDZb motif (**Figures 4H–I**), indicating the necessity of this motif for the Side-IV-Dsyd-1 interaction. In addition, Dsyd-1 accumulated at the membrane where Side-IV had accumulated due to its interaction with Beat-IIb (**Figures 4J–J’’**). Although Side-IV and Liprin-α did not show strong co-localization, their co-localization index nearly doubled when *dsyd-1* was co-transfected in the S2 cells (**Figures 4K–M**). This suggests that Side-IV can form a complex with Liprin-α through Dsyd-1. These findings indicate that Side-IV’s PDZb motif can interact with Dsyd-1 and establish the Dsyd-1/Liprin-α complex crucial for active zone formation.^44, 45^

To validate the notion that Dsyd-1 and Liprin-α function downstream of Side-IV’s cytoplasmic domain, we utilized Side-IV^ΔCyto^, which lacks the ability to transduce intracellular signaling. The knockdown of *dsyd-1* suppressed synaptic induction by Side-IV^FL^, but not Side-IV^ΔCyto^ in R7 (**Figure 4N**), indicating that Dsyd-1 exclusively acts on signaling mediated through the Side-IV’s cytoplasmic domain. Conversely, knockdown of *liprin-α* suppressed synaptic induction by both Side-IV^FL^ and Side-IV^ΔCyto^ (**Figure 4N**), suggesting that Liprin-α affects signaling through both Side-IV’s cytoplasmic domain and the co-receptor(s). Additionally, the knockdown of *kirre* suppressed synaptic induction by Side-IV^ΔCyto^, but not Side-IV^FL^ (**Figure 4N**), indicating that Kirre affects signaling independent of Side-IV’s cytoplasmic domain. Furthermore, Side-IV co-localized with Dsyd-1, Liprin-α, and Kirre in the M1–M3 region of R7 where ectopic synaptic induction occurred (**Figure S5**). These findings demonstrate that Side-IV/Kirre forms a signaling complex with Dsyd-1/Liprin-α through Side-IV’s cytoplasmic domain (**Figure 4O**).

### *side-IV* and *side-VII* are Required for Proximal Synapse Formation in L2 Neurons

By toggled expression experiments of *side-IV* in R7, we identified Side-IV/Beat-IIb as a potential cell-surface protein combination for synaptic induction, albeit in an ectopic context. To investigate their endogenous function, we explored the potential synaptic-partner neurons where Side-IV/Beat-IIb operates using single-cell RNAseq data from the *Drosophila* visual system.^11, 17^ The Side-IV-Beat-IIa/b and Side-VII-Beat-IV interactions were predicted between L2 and L4 neurons **(Figures 5A** and **5B**).^11, 17^ Additionally, endogenous tags of Side-IV exhibited signals in the M2 layer (**Figures S1F** and **S1F’**), indicating that L2 expressed *side-IV*. Among the five subtypes of lamina neurons, only L2 and L4 establish reciprocal synapses specific to the proximal lamina region (**Figures 5A**, **5C**, and **5C’**).^46^ To assess the functional significance of the Beat-Side interactions for synapse formation between L2 and L4 in the proximal lamina, we generated *side-IV* or *side-VII* mutants (see Methods), which were further analyzed (**Figures 5C–H**).

**Figure 5.**
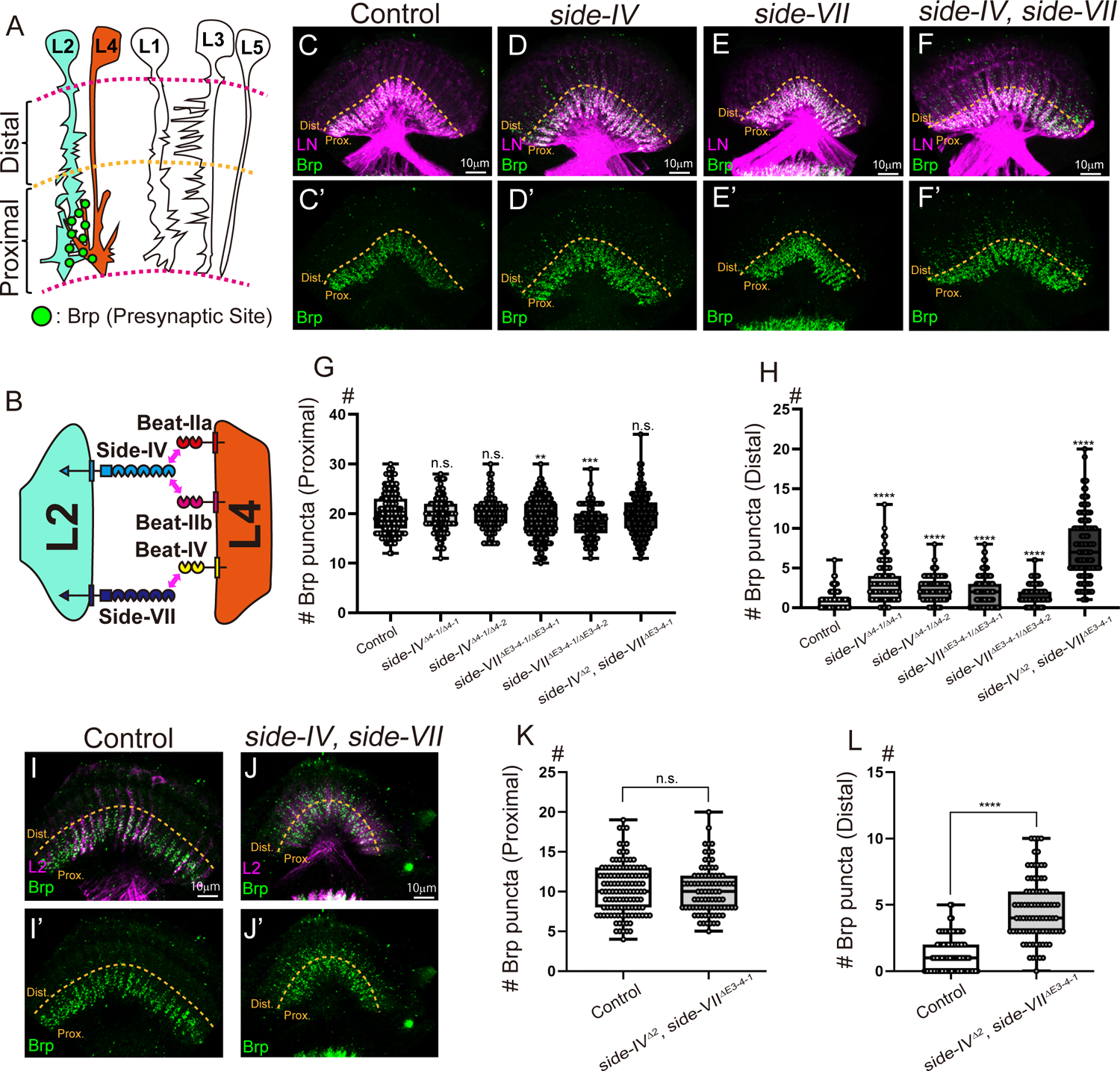
*side-IV* and *side-VII* are Required for Proper Synapse Formation Patterns of the L2 Neurons (A) Scheme of synaptic connection patterns of lamina neurons. (B) Scheme of indicated Beat-Side interactions between L2 and L4 neurons from single-cell RNAseq.^11, 17^ (C–F’) Presynaptic structures and cell morphology of lamina neuron (LN). Control (C and C’), *side-IV^Δ4-1^* mutant (D and D’), *side-VII^ΔE3-4-1^* mutant (E and E’), *side-IV^Δ2^*/*side-VII^ΔE3-4-1^* double mutant (F and F’). Brp puncta (green) of lamina neurons are visualized by combining the STaR method with LN-specific-FLPase (27G05-FLP). Morphology of LN are visualized with LexAop-myrtdTOM (magenta). Dotted lines indicate the boundary between the proximal and distal region. (G) Quantification of the number of synapses in the proximal region (n > 82 axons). (H) Quantification of the number of synapses in the distal region (n > 107 axons). (I–J’) Axonal and synaptic morphology of L2 are visualized with the STaR method by cell-specific FLPase expression in the L2 neurons (53G02-AD (II), 29G11-DBD (III) L2 split-gal4 + UAS-FLP). Control (I and I’), *side-IV^Δ2^*/*side-VII^ΔE3-4-1^*double mutant (J and J’). (K) Quantification of synapses in the proximal region of the L2 neuron (n > 88 axons). (L) Quantification of synapses in the distal region of the L2 neuron (n > 84 axons). Statistical treatment of the number of synapses in the proximal region and distal region were analyzed with the unpaired t-test and Mann-Whitney test, respectively. See also Figures S6 and S7.

Despite Side-IV’s robust ability to organize R7 synapses, the *side-IV* mutant did not exhibit reduced synapse formation in the proximal region (**Figures 5C–D’** and **5G**). This lack of synaptic loss in the absence of synapse-organizing molecules may be attributed to molecular redundancy^47, 48^ such that the disruption of a few genes might be insufficient to decrease synapse numbers. Notably, at least 10 transmembrane interactions were predicted between L2 and L4,^13^ suggesting a high level of molecular redundancy in their synaptic connections. However, the *side-IV* mutant displayed ectopic synapse formation in the distal region (**Figures 5C–D’** and **5H**), resembling the phenotype observed in the *dip-β* mutants.^7^ To elucidate this process further, we conducted rescue experiments with *side-IV* and observed complete rescue of the phenotype with Side-IV^FL^, but only partial rescue of Side-IV^ΔCyto^ (**Figures S6A–D**). This suggests that restoration of the strength of the cell-cell contact alone is insufficient, emphasizing the critical role of cytoplasmic signaling. The *side-VII* mutants also exhibited ectopic synapse formation in the distal region, which was significantly enhanced in the *side-IV*/*side-VII* double mutants (**Figures 5E–H**). Specific knockdown of *side-IV* or *side-VII* in L2 resulted in ectopic synapse formation in the distal region, underscoring the need for *side-IV* and *side-VII* in L2 (**Figures S6E–H**). Furthermore, downregulation of *beat-IIa/b* or *beat-IV* phenocopied the ectopic synapse formation in the distal region (**Figures S6I–P**). To identify the specific lamina neuron responsible for the induction of ectopic synapse formation by disrupting Beat-Side interactions, we analyzed the synapses of each lamina neuron subtype in *side-IV*/*side-VII* double mutant flies. L2 exhibited ectopic synapse formation in the distal region (**Figures 5I–L**), whereas other subtypes did not (**Figure S7**). These findings elucidate the critical role of Side-IV-Beat-IIa/b and Side-VII-Beat-IV interactions in governing proper synapse formation, specifically confined to the proximal region, within L2. Side-IV and Side-VII likely regulate the synaptic specificity of L2 by determining the precise location of synapse formation.

### Side-IV and Side-VII exhibit Beats-Dependent Subcellular Localization to the Proximal Region

To elucidate the roles of Side-IV and Side-VII in determining the L2 synapse formation sites, we investigated their specific localization pattern within the proximal region of the lamina where the presynaptic structures are formed. We generated knock-in strains that incorporated FLPase-based cell-specific tags into *side-IV* and *side-VII* (**Figure S1E**). Remarkably, both Side-IV and Side-VII exhibited distinct and exclusive localization to the proximal region during the pupal stage 72 h after puparium formation (APF) (**Figures 6A–A’’’**). Throughout their development, both *side-IV* and *side-VII* were expressed in L2 neurons, initially showing broad but progressive localization, which later became restricted to the proximal half of the lamina (**Figure S8**). Given the uniform dendritic arbors of the L2 neurons across the distal to proximal regions,^49, 50^ it is plausible that the subcellular localization of Side-IV and Side-VII is modulated in a non-cell autonomous manner, potentially regulated by Beats originating from the L4 neurons.

**Figure 6.**
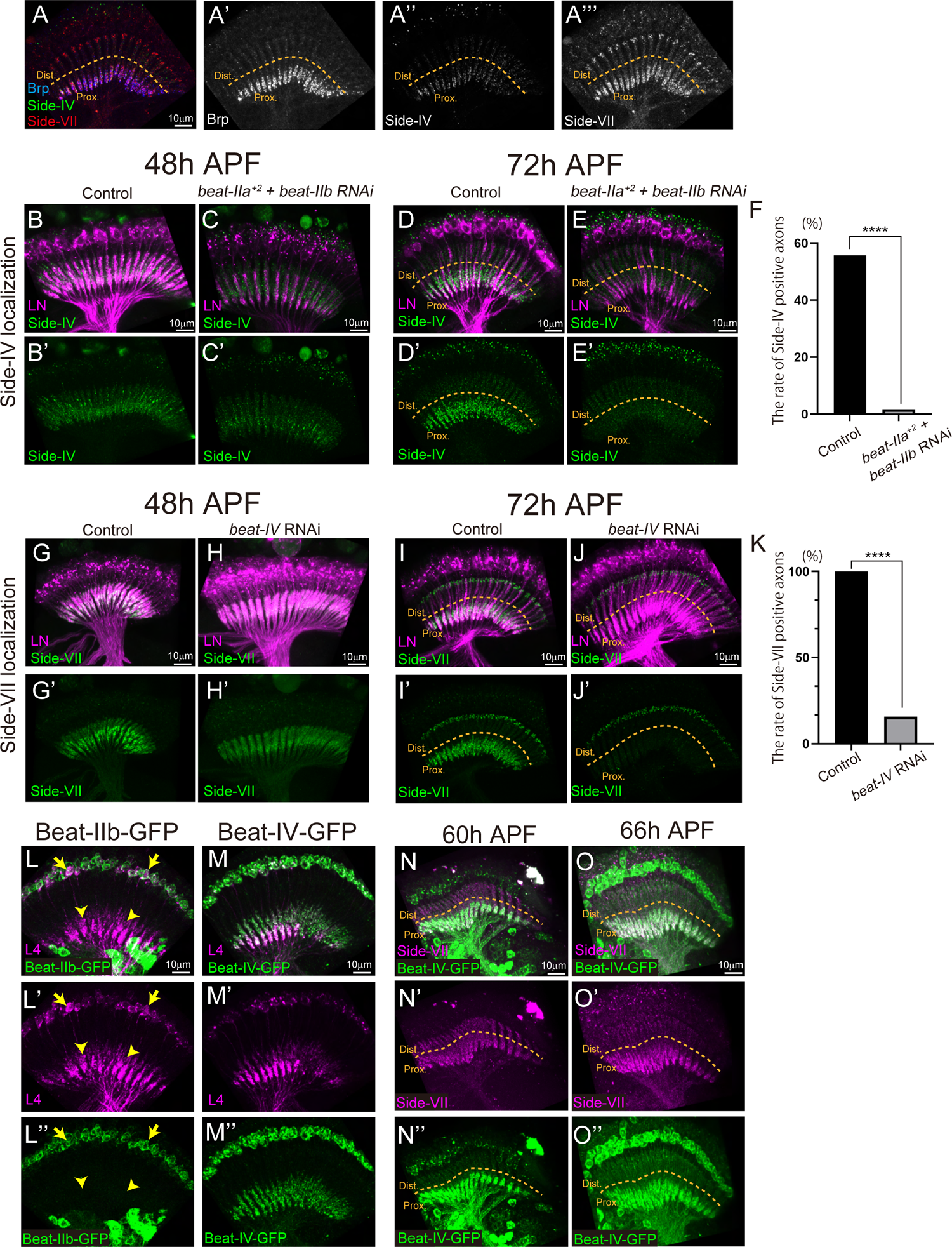
Side-IV and Side-VII Showed Beats-dependent Subcellular Localization to the Proximal Region. (A–A’’’) Mutual localization of endogenous Brp (A’), Side-IV (A’’), and Side-VII (A’’’) at 72 h after pupal formation (APF). *brp*-FsF-V5, *side-IV*-FsF-GFP, and *side-VII*-FsF-mCherry are combined with 27G05-FLP. Dotted lines indicate the proximal and distal boundary. (B–E’) Endogenous localization of Side-IV is visualized by combining *side-IV*-FsF-GFP with 27G05FLP. Control 48 h APF (B and B’), *beat-IIa^+2^*mutant + *beat-IIb* RNAi 48 h APF (C and C’). Control 72 h APF (D and D’), *beat-IIa^+2^* mutant + *beat-IIb* RNAi 72 h APF (E and E’). Side-IV localizations in the proximal region were disrupted by *beat-IIa/b* downregulation. *beat-IIb* are knocked down in lamina neurons (Ay-gal4 + 27G05-FLP) upon *beat-IIa^+2^* mutant. Morphology of LN are visualized with UAS-myr-RFP (magenta). (F) Quantification of the rate of the axons with Side-IV localization in the proximal region at 72 h APF (n > 149 axons). (G–J’) Endogenous localization of Side-VII is visualized by combining *side-VII*-FsF-GFP with 27G05FLP. Control 48 h APF (G and G’), *beat-IV* RNAi 48 h APF (H and H’). Control 72 h APF (I and I’), *beat-IV* RNAi 72 h APF (J and J’). Side-VII localizations in the proximal region were disrupted by *beat-IV* downregulation. (K) Quantification of the rate of the axons with Side-VII localization in the proximal region at 72 h APF (n > 147 axons). (L–L’’) Localization of Beat-IIb-MiMIC-GFP (green) in the lamina neuron at 72 h APF. Morphology of the L4 neuron was visualized with the L4 neuron-specific gal4 driver (GMR31C06-gal4) against UAS-myr-RFP (magenta). No Beat-IIb-GFP signal was observed in the axon of the L4 neuron (arrowheads), although the Beat-IIb-GFP signal was confirmed in the cell body of the L4 neuron (arrows). (M–M’’) Localization of Beat-IV-MiMIC-GFP (green) in the lamina neurons at 72 h APF. (N–O’’) Analysis of the mutual localization of Side-VII and Beat-IV. Side-VII is visualized by combining *side-VII*-FsF-mCherry (magenta) and 27G05FLP. Beat-IV is visualized by *beat-IV*-MiMIC-GFP (green). 60 h APF (N–N’’) and 66 h APF (O–O’’). Statistical treatment of the rate of Side-IV or Side-VII positive axons were analyzed with the Chi-square test. See also **Figure S8**.

We analyzed the localization dependency of Side-IV and Side-VII on Beat-IIa/b and Beat-IV, respectively. With downregulated *beat-IIa*/*b*, Side-IV failed to localize to the proximal region at 72 h APF (**Figures 6B–F**). Similar to the impact of *beat-IIa/b* on Side-IV’s localization, the downregulation of *beat-IV* disrupted Side-VII’s localization 72 h APF (**Figures 6G–K**). These findings indicated that the subcellular localization of Side-IV and Side-VII are Beat-IIa/b and Beat-IV-dependent, respectively.

We further examined the localization of Beat-IIb and Beat-IV using MiMIC-GFP transgenics.^38^ We primarily focused on Beat-IV, since Beat-IIb-MiMIC-GFP exhibited localization only in cell bodies (**Figures 6L–L’’**, yellow arrows), possibly due to defects in transport to L4 dendrites. Beat-IV-MiMIC-GFP localization was observed in the proximal region prior to Side-VII localization (60 APF) (**Figures 6M–O’’**). These results suggest that Beat-IV localizes to the proximal region before Side-VII, indicating that it plays a role in directing the subcellular localization of Side-VII.

### Complex Formation of Side-IV with Kirre and Dysd-1 Restricts Loci of Synapse Formation and Inhibits Miswiring

Our analyses of the function and localization of Side-IV and Side-VII provide compelling evidence for their role in directing synapse formation within the proximal region of L2. However, there is an intriguing question regarding the occurrence of ectopic synapse formation in the distal region, as observed in the *side-IV* or *side-VII* mutants. Previous studies have demonstrated that *dip-β* mutants exhibit miswiring to alternative neurons, resulting in ectopic synapse formation in the distal region, and that DIP-β directs L4 neurons to favor L2 as a synaptic partner over other neurons.^7^ This suggests that both Side-IV and Side-VII may encourage L2 to select L4 while inhibiting miswiring to alternative neurons. Consequently, the absence of either Side-IV or Side-VII could highlight the presence of molecules that promote the miswiring of L2 to alternative neurons.

Our analysis of *side-IV* in R7 neurons revealed that Kirre can function as a co-receptor for Side-IV (**Figure 3**). Previous research has shown that the *kirre* mutants result in a complete loss of the postsynaptic structures of L4,^51^ indicating that Kirre is a major presynaptic regulator in L2. Furthermore, single-cell-RNAseq data predicted homophilic self-interactions of Kirre and heterophilic Kirre-Rst interactions, between L2 and all other subtypes of lamina neurons.^13, 16^ Based on these findings, we hypothesized that Kirre cooperates with Side-IV to establish connectivity between L2 and L4 but, in the absence of Side-IV, Kirre could induce synaptic connections indiscriminately. A protein localization analysis revealed that Kirre exhibits robust localization in the proximal region, similar to Side-IV (**Figures 7A–B**). However, Kirre displayed broader localization than Side-IV, and some punctate signals were observed in the distal region (**Figures 7A’’’**, yellow arrowheads, **7B**, and **7C**), indicating its positive interaction with Kirre/Rst in both regions. To investigate whether Kirre is responsible for the ectopic synapse formation in the *side-IV* mutants, we examined genetic interactions between *kirre* mutants^52^ and *side-IV* or *side-IV/side-VII* double mutants (**Figures 7D–K**). The deletion of one copy of *kirre* resulted in a reduction in the number of synapses in the proximal region (**Figure 7J**) and, importantly, the suppression of ectopic synapse formation in the distal region (**Figure 7K**). These data indicate that Kirre promotes the miswiring of L2 with alternative neurons in the distal region in the absence of *side-IV* and/or *side-VII*.

**Figure. 7.**
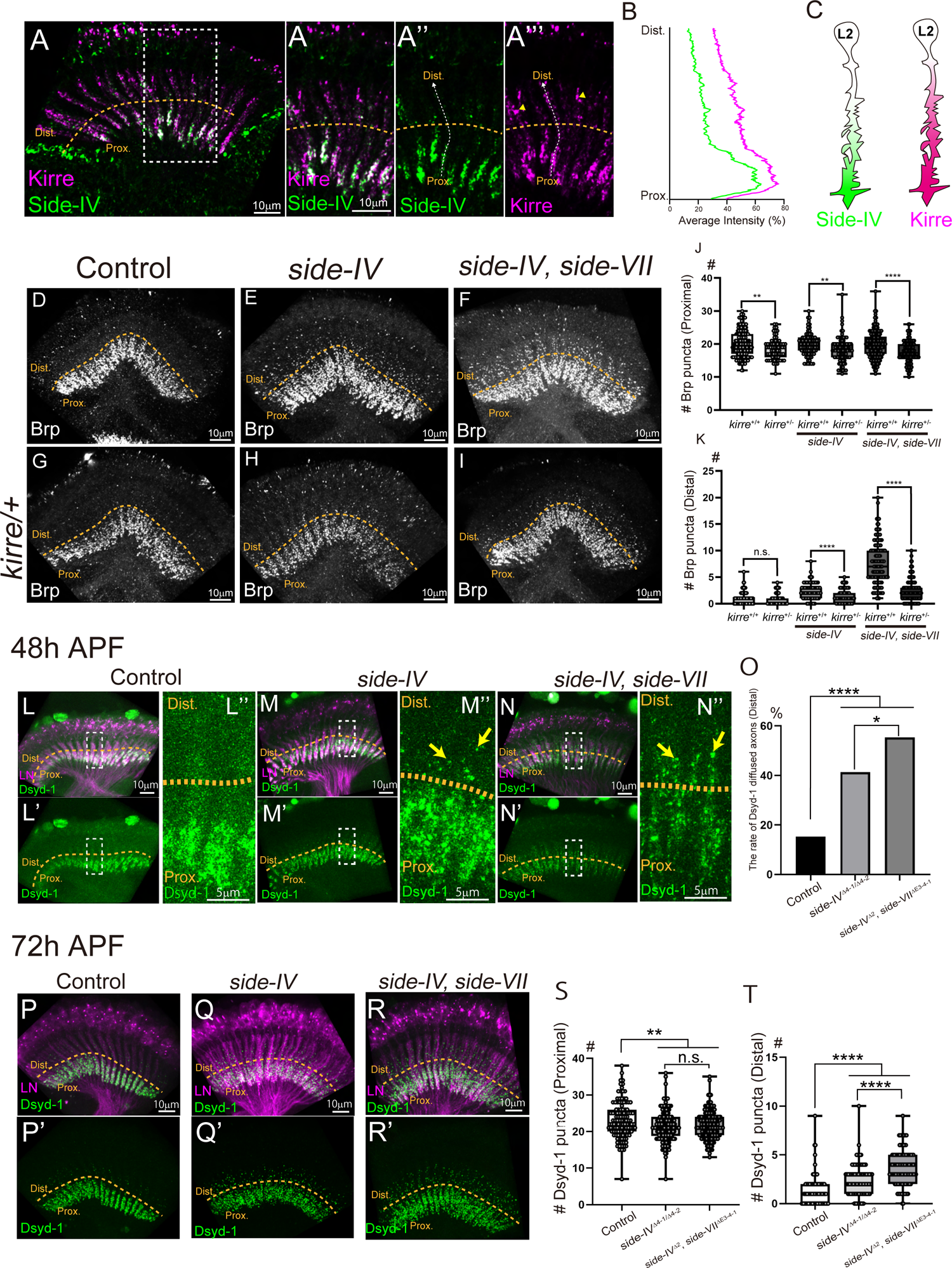
Side-IV Limit the Dysd-1 Localization in the Proximal and Inhibit the Synapse Formation by Kirre in the Distal Region. (A) Mutual localization Kirre and Side-IV on L2 neuron at 72 h APF. Merge (A), magnified Merge (A’), Side-IV-GFP (A’’), Kirre-HA (A’’’). Compared with Side-IV localization, Kirre puncta signals were observed in the distal region (arrowheads in A’’’). UAS-*side-IV*-GFP and UAS-*kirre*-HA are combined with L2-specific-gal4 (GMR16H03-gal4). Dotted lines indicate the proximal and distal boundary. (B) Fluorescent intensity of Side-IV-GFP (green) and Kirre-HA (magenta) measured from proximal to distal across L2 axons as shown in A’’ and A’’’ (dotted lines). Average intensity of 29 axons was calculated. (C) Schematic of Side-IV (green) and Kirre (magenta) localization in the L2 neuron. (D–I) Presynaptic structures of lamina neurons visualized by STaR method. Control (D, same image as **5C’**), *side-IV^Δ4-1/ Δ4-2^*mutant (E), *side-IV^Δ2^*/*side-VII^ΔE3-4-1^*double mutant (F, same image as **5F’**), *kirre^sps^* heterozygous mutant (G), *kirre^sps^* heterozygous and *side-IV^Δ4-1/ Δ4-2^*mutant (H), *kirre^sps^* heterozygous and *side-IV^Δ2^*/*side-VII^ΔE3-4-1^* double mutant (I). Dotted lines indicate the proximal and distal boundary. (J) Quantification of synapses in the proximal region (n > 69 axons). (K) Quantification of synapses in the distal region (n > 79 axons). (L–N’’) Localization of endogenous Dsyd-1 in lamina neurons is visualized combining *dsyd-1*-FsF-GFP with 27G05FLP at 48 h APF. Control (L–L’’), *side-IV^Δ4-1/ Δ4-2^*mutant (M–M’’), *side-IV^Δ2^*/*side-VII^ΔE3-4-1^* double mutant (N–N’’). Arrows indicate diffused Dsyd-1 signal to the distal region. (O) Rate of Dsyd-1 diffused axons in the distal region at 48 h APF (n > 92 axons). (P–R’) Localization of Dsyd-1 in lamina neuron at 72 h APF Control (P and P’), *side-IV^Δ4-1/ Δ4-2^*mutant (Q and Q’), *side-IV^Δ2^*/*side-VII^ΔE3-4-1^*double mutant (R and R’). (S) Quantification of the number of Dsyd-1 puncta in the proximal region (n > 120 axons). (T) Quantification of the number of Dsyd-1 puncta in the distal region (n > 96 axons). Statistical treatment of the number of Brp or Dsyd-1 puncta in the proximal region and distal region were analyzed with the unpaired t-test and Mann-Whitney test, respectively. Rate of Dsyd-1 diffused axon was analyzed with by the Chi-square test.

The mechanism through which Side-IV exerts its inhibitory effect on ectopic synapse formation in the distal region of the wild type remains elusive. One potential explanation is the presence of competitive mechanisms that prevent ectopic synapse formation. The specific localization of Side-IV and Side-VII to the proximal region (**Figure 6**) suggests that they may compete with molecules operating in the distal region. Competitive synapse formation has previously been documented in R7 neurons and is characterized by the extension of multiple filopodia for the exploration of potential synaptic partners. At any given time point, only a limited number of these filopodia successfully recruit synapse-seeding factors and stabilize.^3^ Consequently, synaptic loci compete for the available pool of synapse-seeding factors, which may include Dsyd-1 and Liprin-α.^3^ To investigate the competitive role of Dsyd-1, we employed a FLPase-based cell-specific GFP-tag knock-in at the Dsyd-1 locus and examined its endogenous localization (**Figures 7L–T**) (see Methods). In the control, Dsyd-1 localized to the proximal region 48 h APF (**Figures 7L–L’’**) and its signals became punctate but remained active in the proximal region 72 h APF (**Figures 7P** and **7P’**). However, both *side-IV* and the *side-IV/side-VII* double mutant flies exhibited leakage of Dsyd-1 signals in the distal region as early as 48 h APF (**Figures 7M– N’’’**, yellow arrows, and **7O**). The number of Dsyd-1 puncta decreased in the proximal region and increased in the distal region in the side mutants 72 h APF (**Figures 7Q–T**). These findings suggest that Side-IV is responsible for anchoring Dsyd-1 localization to the proximal region, preventing opportunities for synapse formation by molecules distributed across the distal region. Taken together, our results indicate that Side-IV collaborates with Kirre and Dsyd-1 to establish L2-L4 synapse specificity in the proximal region, thereby restricting synapse formation loci and inhibiting their miswiring.

## DISCUSSION

This study identified Side-IV/Beat-IIb as a novel combination of cell-surface recognition molecules that can induce synapse formation through establishing a cluster complex and signaling via its co-receptor Kirre or cytoplasmic signaling involving Dsyd-1/Liprin-α in the *Drosophila* R7 photoreceptor. Also, we identified the regulatory function of Side-IV in determining the sites of L2 synapse formation by anchoring Dsyd-1 to prevent miswiring, with the subcellular localization controlled by Beat-IIa/b. Consequently, we propose a model of cell-surface protein complex formation between the Beat and Side families that may play a crucial role in establishing synaptic specificity (**Figure 8**).

**Figure. 8.**
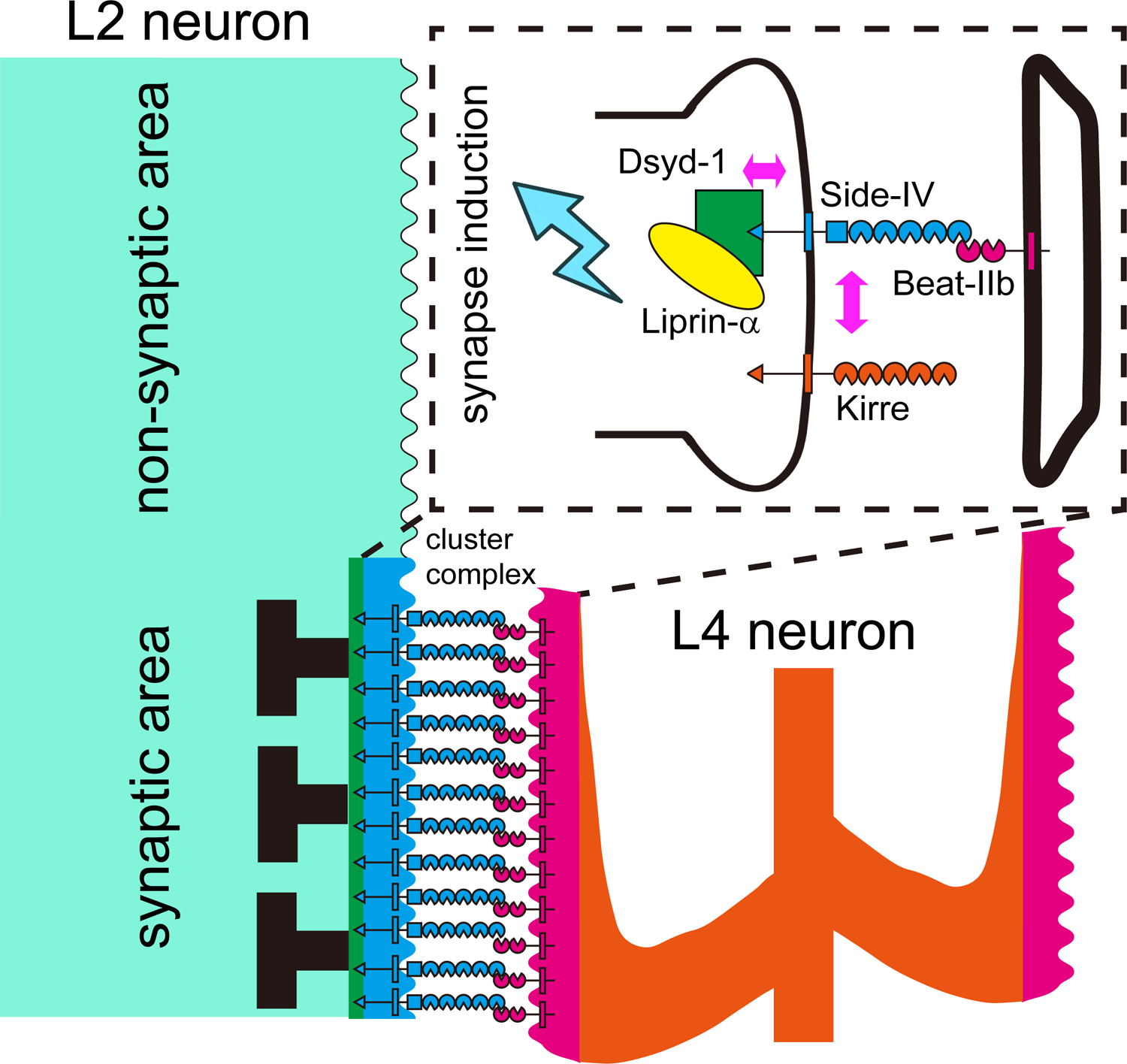
Side-IV-Beat-IIb’s Complex Formation Model. New cell-surface recognition molecule combination Side-IV/Beat-IIb made cluster complex and transduced bifurcated signaling through co-receptor, Kirre and scaffold protein complex, Dsyd-1-Liprin-α. Side-IV can specific the loci of synapse formation and anchor the Dsyd-1 to inhibit miswiring.

### Side-IV Establishes a Signaling Complex with Kirre and Dsyd-1

An interesting finding was that Side-IV can induce synapse formation even in the absence of its cytoplasmic domain (**Figures 2E–E’’’** and **2J**). This led us to hypothesize that Side-IV may exert its signaling function through co-receptors. We have demonstrated that Side-IV and its co-receptor Kirre synergistically mediate signal transduction via their extracellular domains. Kirre, a well-known member of the irre cell recognition module (IRM) family,^41^ has diverse functions ranging from nervous system-related processes such as axon guidance and synapse formation to myoblast aggregation and fusion, as well as cell sorting in diverse biological contexts.^41, 51, 52^ It was intriguing to discover the *cis*-regulation of Side-IV by the well-known cell adhesion molecule. Notably, co-expression of *side-IV* and the other IRM members, *hbs* and *sns,* did not enhance the phenotype (data not shown), indicating that Kirre may function as a unique co-receptor for Side-IV. Interestingly, our findings also suggested the presence of signaling redundancies in the Side-IV/Kirre pathway. Our co-overexpression experiments in R7 demonstrated that, if the signaling pathway of either Side-IV or Kirre was active, as indicated by the presence of their respective cytoplasmic domains, the complex could induce a more pronounced phenotype (**Figures 3G–P**). This study indicates that individual Beat-Side family proteins may possess unique co-receptors with cytoplasmic redundancy that interact to generate diverse neuronal connections.

Furthermore, our study suggests that Dsyd-1 and Liprin-α work downstream of Side-IV. Strong co-localization between Side-IV and Dsyd-1 was observed but this significantly decreased upon deletion of the Side-IV PDZb motif (**Figures 4G–I**). Notably, the knockdown of *dsyd-1* specifically impeded synaptic induction by cytoplasmic signaling of Side-IV (**Figure 4N**). Additionally, Liprin-α co-localized with Side-IV through its interaction with Dsyd-1 and the knockdown of *liprin-a* caused considerable suppression of synaptic induction by Side-IV (**Figures 4J–M**). These findings suggest that Dsyd-1 interacts with the Side-IV PDZb motif and collaborates with Liprin-α to co-contribute to synapse formation. It is worth mentioning that in *Drosophila,* other synapse organization molecules Nrx-1 and Lar interact with Dysd-1 and Liprin-α, respectively.^44, 53^ Consequently, Side-IV may compete with these molecules to recruit Dsyd-1 or Liprin-α, thereby exerting its influence on the synaptic preferences of the neuron.

### Subcellular Localization of Side-IV and Side-VII Establishes L2-L4 Connectivity

Although Side-IV demonstrates robust synapse induction abilities in R7, the *side-IV* mutant did not exhibit any decrease in the number of synapses in lamina neurons (**Figures 5C–D’**, and **G**). This observation is quite common with other synapse organizers, such as Neurexins and Neuroligins; even knockdown or mutation of their three paralogs only marginally affects synapse numbers despite their potent synapse-organizing capabilities.^47, 48^ Conversely, a notable increase in ectopic synapse formation was observed in the distal region of the *side-IV* mutant (**Figures 5C–D’** and **5H**).

Based on the expression data, three Beat-Side interactions were predicted between L2 and L4 (**Figures 5A** and **5B**),^11, 17^ and the disruption of these interactions was significantly correlated with an elevation in the number of ectopic synapses induced in the distal region. Notably, ectopic synapses were found exclusively in L2 and not in other subtypes of lamina neurons (**Figures 5I–L** and **S7**). These findings indicate that multiple members of the Beat and Side families redundantly contribute to the determination of synapse formation sites in L2, particularly within the proximal region. This was supported by our subcellular localization analysis, which revealed distinct distribution patterns of Side-IV and Side-VII to the proximal region in L2. These were regulated by Beat-IIa/b and Beat-IV, respectively (**Figures 6B–K**). Despite L2 exhibiting uniform dendritic arborization from the proximal to the distal regions,^49, 50^ the interplay between Beat-Side interactions and the nonuniform cellular morphology of L4 (with dendrite distribution to the proximal region)^49^ appears to establish nonuniform synapse formation restricted to the proximal region of L2. Intriguingly, the DIP-β protein operating at the loci of L4 synapse formation also exhibited significant accumulation in the proximal region at 72 h APF,^7^ resembling the localization patterns of Side-IV and Side-VII. Therefore, the collaboration between DIP-Dpr and Beat-Side families likely plays a crucial role in establishing L2-L4 connectivity.

### Complex Formation of Side-IV Links Synaptic Specificity and Synaptogenesis

Synaptogenesis possesses a dual nature, characterized by the robustness of its neural circuit establishment and its potential for indiscriminate connections.^4,5,7,8^ Molecular coding, such as that for the recognition of cell-surface molecules, likely plays a vital role in determining synaptic partners.^2,6^ However, the loss of such molecules can sometimes result in miswiring with alternative partners.^7,8^ Neurons have the ability to form synapses with various partners, and cell-surface molecules can create preferences for certain synaptic partners. Initially, we thought these synaptic partner preferences were determined by the interaction affinities of cell-surface recognition molecules. However, increasing the affinity between R7 and postsynaptic partners through homophilic interactions of CadE or hSDK2 alone did not induce synapse formation (**Figure S3**). Moreover, the incomplete rescue of the ectopic synapse formation in *side-IV* mutants by Side-IV^ΔCyto^ indicated that intracellular signaling mechanisms were necessary to enhance preferences and facilitate synaptic connection formation (**Figures S6A– D**).^54^

Previous studies have elucidated the role of DIP-β in facilitating the preferential formation of synapses between L4 and L2 neurons.^7^ However, the exact intracellular signaling mechanisms of DIP-β remain elusive due to the absence of apparent intracellular signaling motifs in DIP-β.^16, 55^ The phenotypic similarities between *dip-β* and *side-IV* mutants suggest that Side-IV also plays a role in facilitating L2 neurons’ preference to synapse formation with L4 neurons. Our study demonstrated that Dsyd-1 and Liprin-α, important signaling factors for active zone formation,^44, 45^ operate downstream of Side-IV. Furthermore, Side-IV can anchor the localization of Dsyd-1 to the proximal region, resulting in the accumulation of Dsyd-1 at the L2-L4 contact sites and increased preference of L2 for L4 over other neurons for synapse formation. This competition for the synapse-seeding factor Dsyd-1, as a limited resource, ensures relative preference for synapse formation,^3,56^ thereby inhibiting miswiring of L2 neurons. Similarly, Kirre, another major driver of L2-L4 synaptogenesis,^51^ can promote miswiring if not properly regulated (**Figures 7D–K**). These findings underscore the importance of signaling complex establishment between cell-surface molecules and synapse formation factors, with the appropriate ratio and subcellular localization, in achieving synaptic specificity. In this study, we have presented an example of such complex formation through the interaction of Side-IV, Kirre, and Dsyd-1, which initiates synaptogenesis, coupled with the *trans-*synaptic interaction between Side-IV and Beat-IIb, which determines synaptic specificity.

## ACKNOWLEDGMENTS

We gratefully acknowledge Dr. Filipe Pinto-Teixeira at Molecular, Cellular and Developmental biology unit (MCD) in Center for Integrative Biology for providing the side-IV[MI10049]-T2AGal4 flies. We thank the Bloomington Drosophila Stock Center, Vienna Drosophila Resource Center (VDRC), Kyoto Drosophila Stock Center (Kyoto-Fly), and FlyORF for providing fly stocks. We thank Enago (www.enago.jp) for the English language review. This work was supported by JSPS KAKENHI #21J12660 (J. O.), a Grant-in-Aid for Scientific Research (B) and a Grant-in-Aid for Scientific Research on Innovative Areas from MEXT (#16H06457, #21H05682, #21H02483, and #23H04220 (T. S.)), a Takeda Visionary Research Grant from the Takeda Science Foundation (T. S.).

---

**Figure S1.**
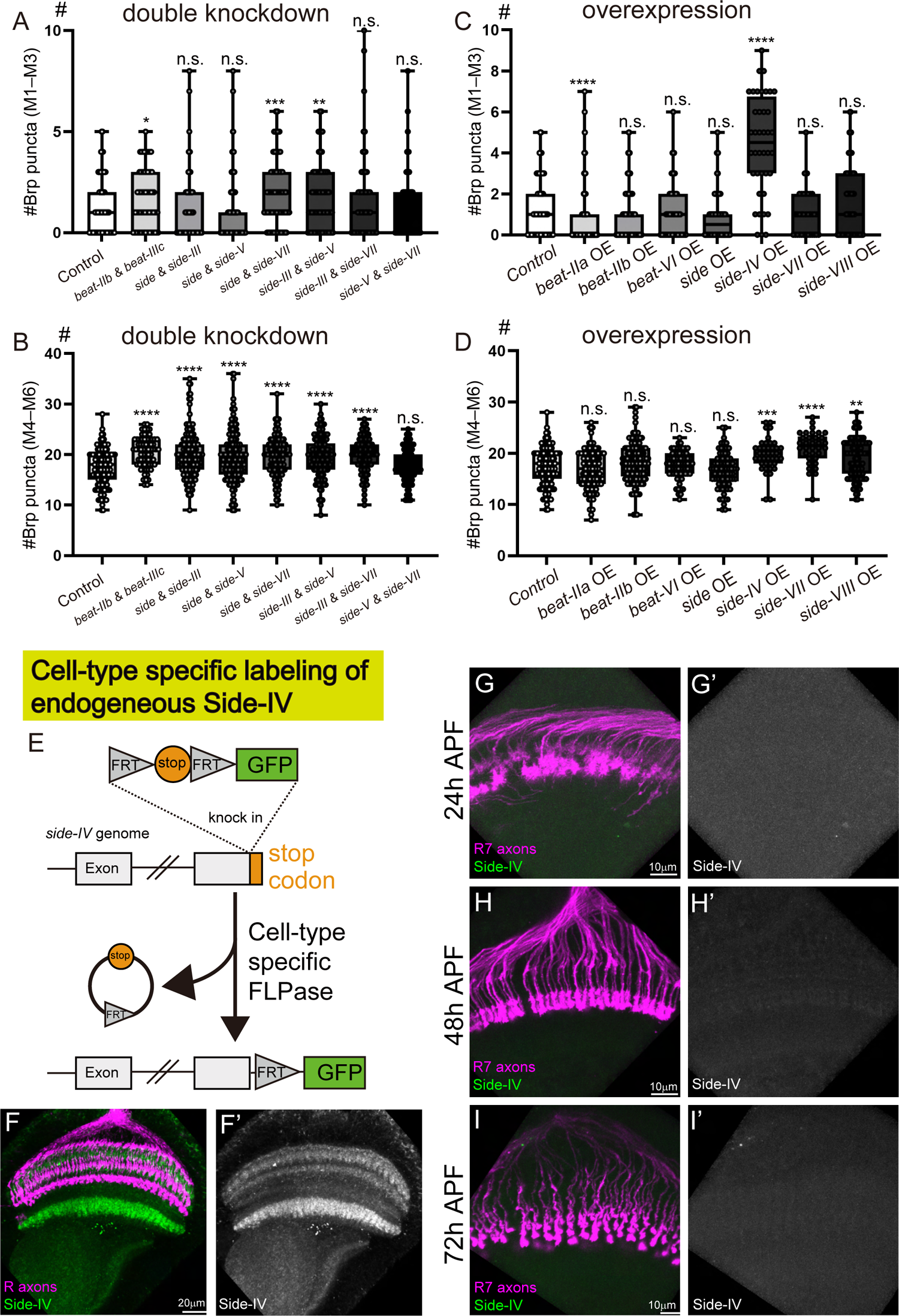
Mis-expression of *side-IV* in R7 induces ectopic synapse formation in the M1–M3 layer (related to Figure 1) (A–D) Beat/Side family members were either knocked down or overexpressed by GMR-gal4. R7 synapses are visualized by combining STaR technique with 20C11-FLP and quantified. (A and B) Beat/Side double knockdown experiments. Quantification of the number of synapses in the M1–M3 layer (A) and M4–6 layer (B) per axon (n 96 axons). (C and D) Beat/Side overexpression (OE) experiments. Quantification of the number of synapses in the M1–M3 layer (C) and M4–6 layer (D) per axon (n > 34 axons). (E) Scheme of tissue-specific visualization of endogenous Side-IV. The FRT-stop-FRT-GFP cassette was knocked in just before the stop codon of *side-IV*. By cell specific expression of FLPase, endogenous localization of Side-IV are visualized in a cell-specific manner. (F and F’) Localization of Side-IV in the optic lobe at adult stage. heatshock-FLP was used to visualize Side-IV (green) in all cells. R axons are stained by mAb24B10 (magenta). (G–I’) Expression of endogenous *side-IV* in R7 at developmental stages. Endogenous Side-IV in R7 is visualized combining *side-IV*-FsF-GFP with 20C11-FLP. R7 axons are visualized combining UAS-myr-RFP with R7-specific-gal4 (magenta, GMR-FsF-gal4 + 20C11-FLP). 24 h APF (G and G’), 48 h APF (H and H’), 72 h APF (I and I’). No Side-IV-GFP signals were observed at any stages, indicating that R7 does not endogenously express *side-IV*. Statistical treatment of the number of the number of synapses in the M1–M3 layer and M4–M6 layer were analyzed with the Mann-Whitney test and unpaired t-test, respectively. In this and all following statistical treatment; n.s. *P* > 0.05, * *P* ≤ 0.05, ** *P* ≤ 0.01, *** *P* ≤ 0.001, **** *P* ≤ 0.0001.

**Figure S2.**
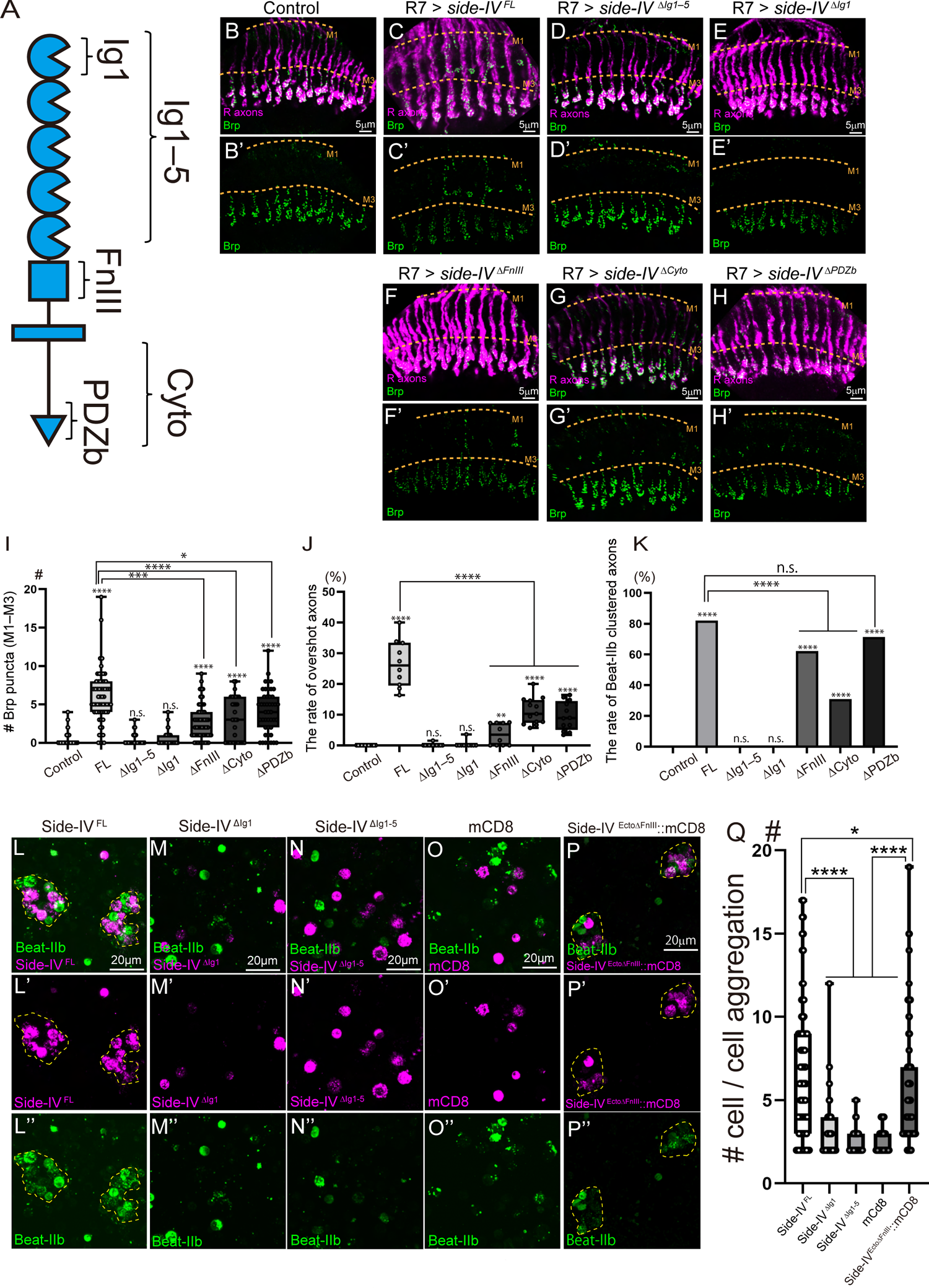
Ig domain of Side-IV is necessary for *trans*-interaction with Beat-IIb that is crucial for synapse induction (related to Figure 2) (A) Domain structure of Side-IV. (B–H’) Brp-GFP signal of R7 with overexpression of *side-IV* domain deletion variants by R7-specific-gal4 (GMR-FsF-gal4 + 20C11-FLP). Brp-GFP puncta (green) in R7 are visualized by combining the STaR method with 20C11-FLP. R axons are stained by mAb24B10 (magenta). Control (B and B’), R7 > *side-IV ^FL^* (C and C’, same images as 1C and C’), R7 > *side-IV ^ΔIg1-5^*(D and D’), R7 > *side-IV ^ΔIg1^*(E and E’), R7 > *side-IV ^ΔFnIII^*(F and F’), R7 > *side-IV ^ΔCyto^*(G and G’), R7 > *side-IV ^ΔPDZb^*(H and H’). Dotted lines indicate region of M1–M3 layer. (I) Quantification of the number of synapses in the M1–M3 layer per axon (n > 33 axons). (J) Quantification of the rate of overshot axons per optic lobe (n > 10 axons). (K) Quantification of the rate of Beat-IIb clustered axons (n > 68 axons). (L–P’’) Cell aggregation assay with *Drosophila* S2 cell. S2 cells were individually transfected with UAS-*side-IV-* mCherry variants or UAS-*mcd8*-mCherry or UAS-*beat-IIb*-GFP together with Actin5C-gal4. After the transfection, cells were subjected to aggregation assay. Side-IV ^FL^-mCherry vs Beat-IIb-GFP (L–L’’), Side-IV ^ΔIg1^-mCherry vs Beat-IIb-GFP (M–M’’), Side-IV ^ΔIg1–5–mCherry^ vs Beat-IIb-GFP (N–N’’), mCd8-mCherry vs Beat-IIb-GFP (O–O’’), and Side-IV ^EctoΔFnIII^::mCD8 vs Beat-IIb-GFP (P–P’’). Dotted lines represent cell aggregations. (Q) Quantification of the number of cell per cell aggregation (n > 40 cell aggregations). Statistical treatment of the number of synapses in the M1–M3 layer, the rate of overshoot axons, and the number of cell per cell aggregation were analyzed with the Mann-Whitney test. Statistical treatment of the rate of Beat-IIb clustered axons was performed with the Chi-square test.

**Figure S3.**
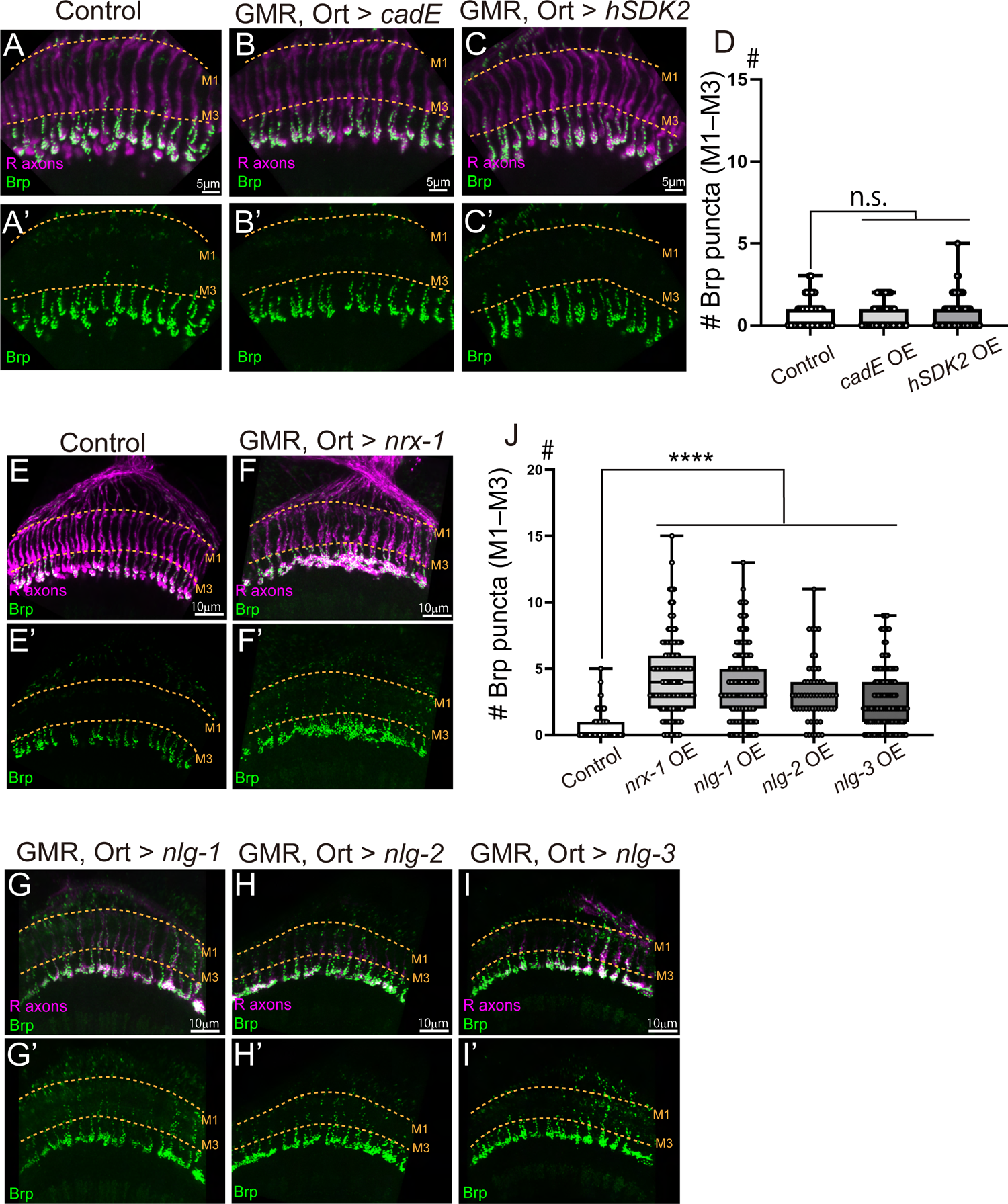
Enhancing physical affinity alone between neurons are insufficient for synapse induction. (related to Figure 2) (A–D) Brp-GFP signal of R7 with overexpression (OE) of the molecules which have homophilic interaction in R7 and postsynaptic neurons by combining GMR-gal4 and Ort-gal4. Brp-GFP puncta (green) in R7 is visualized by combining the STaR technique with 20C11-FLP. R axons are stained by mAb24B10 (magenta). Control (A and A’), *cadE* OE (B and B’), *hSDK2* OE (C and C’). Dotted lines indicate region of M1–M3 layer. (D) Quantification of the number of synapses in the M1–M3 layer per axon (n > 67 axons). (E–I’) Brp-GFP signal of R7 with OE of *nrx-1* or *nlg1–3* with FlySAM in both R7 and postsynaptic neuron by combining GMR-gal4 and Ort-gal4. Control (E and E’), *nrx-1* OE (F and F’), *nlg-1* OE (G and G’), *nlg-2* OE (H and H’), *nlg-3* OE (I and I’). (I) Quantification of the number of synapses in the M1–M3 layer per axon (n > 58 axons). Statistical treatment of the number of synapses in the M1–M3 layer was analyzed with the Mann-Whitney test.

**Figure S4.**
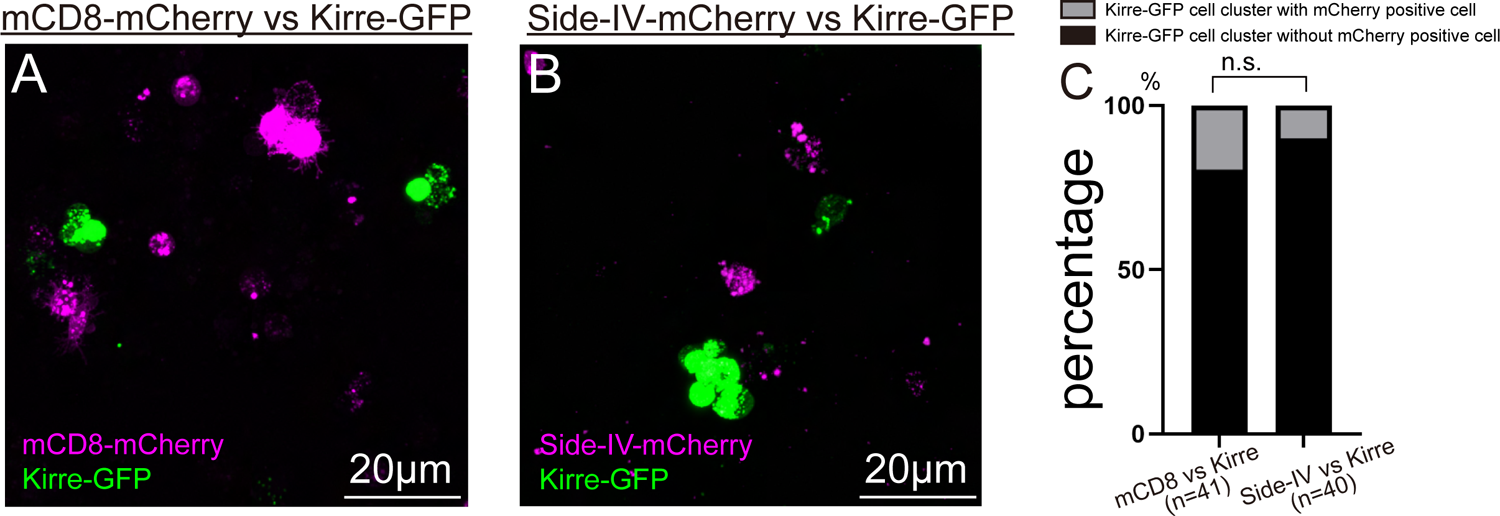
Side-IV does not interact with Kirre in *trans* (related to Figure 3) (A) Aggregation assay of S2 cell transfected Actin5C-gal4 and UAS-*mcd8*-mCherry against Actin5C-gal4 and UAS-*kirre*-GFP. Homophilic interaction of Kirre causes Kirre-transfected S2 cells to form aggregates (Kirre-GFP cluster). (B) Aggregation assay of S2 cell transfected Actin5C-gal4 and UAS-*side-IV*-mCherry against Actin5C-gal4 and UAS-*kirre*-GFP. (C) Quantification of the rate of Kirre-GFP cluster that contains mCherry positive cell. If Side-IV interacts with Kirre in *trans*, more Kirre-GFP clusters should contain *side-IV*-mCherry expressing cells than *mcd8*-mCherry expressing cells. There was no significance of the rate of Kirre-GFP cluster that has mCherry positive cell between aggregation assay of mCd8-mCherry vs Kirre-GFP and Side-IV-mCherry vs Kirre-GFP, indicating Side-IV did not interact with Kirre in *trans*. The sample size is indicated (n). Statistical treatment of the rate of the Kirre-GFP cell cluster with mCherry positive cell was performed with the Chi-square test.

**Figure S5.**
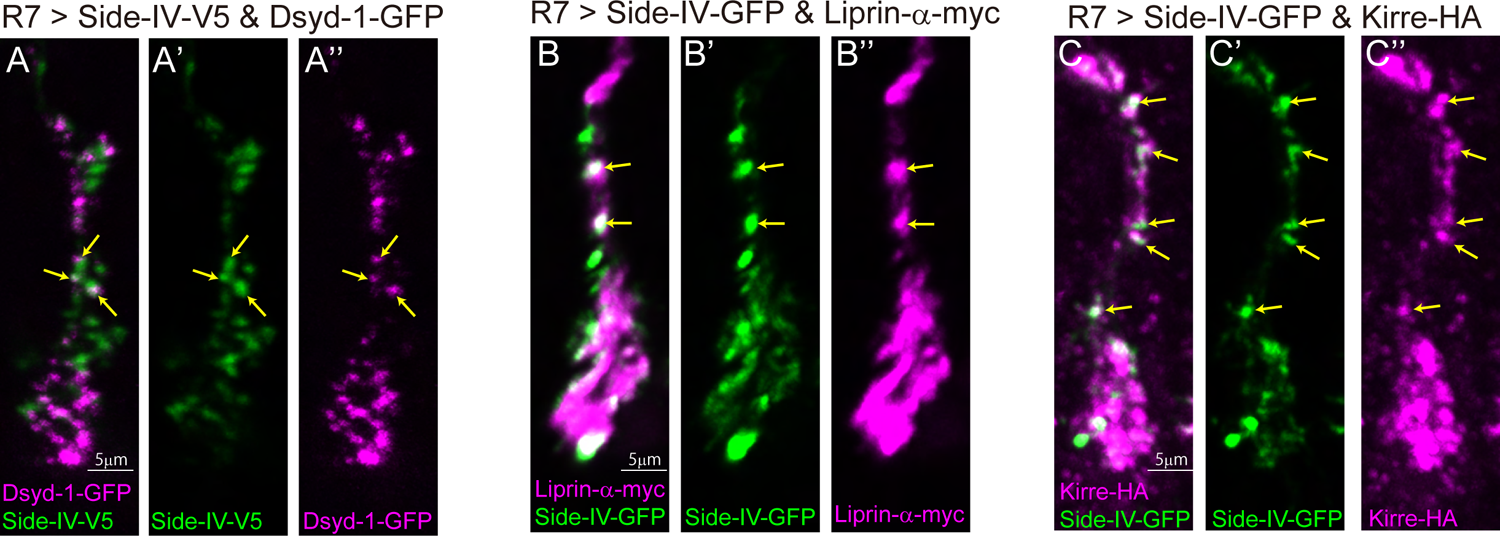
Side-IV co-localized with Dsyd-1, Liprin-α, and Kirre around the M1-M3 layer during development. (related to Figure 4) (A–C’’) Mutual localization of Side-IV and Dsyd-1, Liprin-α, or Kirre in R7 at 48h APF. Arrows indicate the co-localizations around the M1–M3 layer. Endogenous localization of Dsyd-1 is visualized by combining *dsyd-1*-FsF-GFP with 20C11-FLP. Localization of Side-IV, Liprin-α, and Kirre are visualized by expressing tagged proteins by R7-specific-gal4 (GMR-FsF-gal4 + 20C11-FLP). Side-IV-V5 and Dsyd-1-GFP merge (A), Side-IV-V5 (A’), Dsyd-1-GFP (A’’). Side-IV-GFP and Liprin-*<ι>α</i>*-myc merge (B), Side-IV-GFP (B’), Liprin-α-myc (B’’). Side-IV-GFP and Kirre-HA merge (C), Side-IV-GFP (C’), Kirre-HA (C’’).

**Figure S6.**
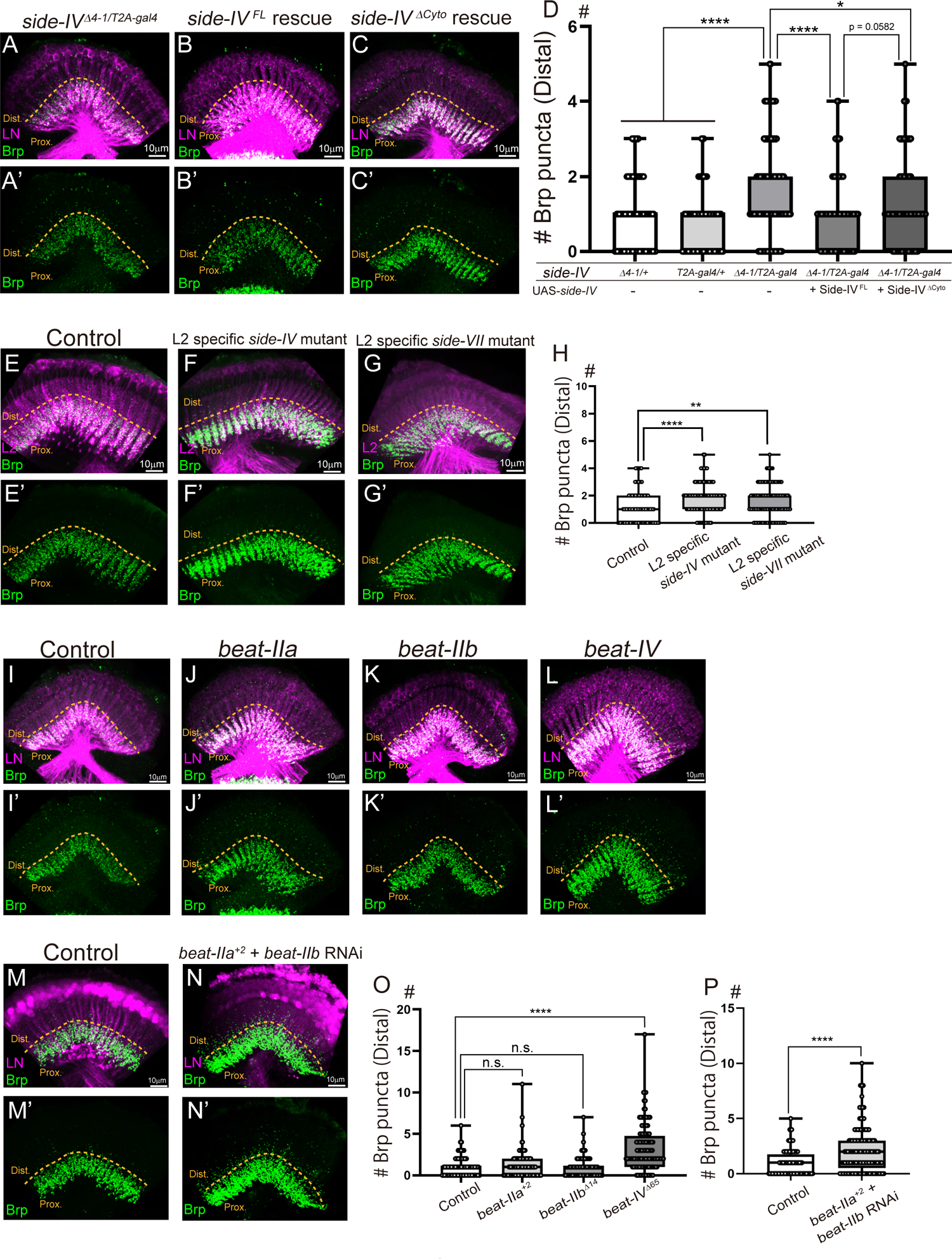
Disruption of Side-IV-Beat-IIa/b or Side-VII-Beat-IV interactions results in ectopic synapse formation in the distal region (related to Figure 5) (A–C’) Presynaptic structures and cell morphology of Lamina neuron (LN). *side-IV^Δ4-1/T2A-gal4^*(A and A’), *side-IV^Δ4-1/T2A-gal4^*+ Side-IV ^FL^ (B and B’), *side-IV^Δ4-1/T2A-gal4^*+ Side-IV *^<ι>Δ</i>^*^Cyto^ (C and C’). Brp puncta (green) of lamina neurons are visualized by combining the STaR technique with LN-specific-FLPase (27G05-FLP). Morphology of LN are visualized with LexAop-myrtdTOM (magenta). Dotted lines indicate the boundary between the proximal and distal region. (D) Quantification of the number of synapses in the distal region in the rescue experiment (n > 88 axons). (E–G’) Brp puncta of lamina neuron in L2 specific *side-IV* or *side-VII* mutant. Morphology of L2 neuron (magenta) are visualized by combining UAS-myr-RFP with L2-specific gal4 driver (GMR16H03-gal4). L2-specific *side-IV* or *side-VII* mutant means *side-IV^Δ4-1^*or *side-VII^ΔE3-4-1^*heterozygote with L2-specific RNAi (GMR16H03-gal4, UAS-RNAi). Control (E and E’), L2 specific *side-IV* mutant (F and F’), L2 specific *side-VII* mutant (G and G’). Orange dotted lines indicate the boundary between the proximal and distal region. (H) Quantification of the number of synapses in the distal region (n > 96 axons). (I–L’) Brp puncta of lamina neuron in *beat* mutants. Control (I and I’), *beat-IIa^+2^* mutant (J and J’), *beat-IIb^Δ14^* mutant (K and K’), *beat-IV^Δ65^* mutant (L and L’). (M–N’) Brp puncta of lamina neurons when *beat-IIb* were knocked down on *beat-IIa* mutant background. Control (M and M’), *beat-IIa^+2^* mutant + *beat-IIb* RNAi (N and N’). *beat-IIb* was knocked down specifically in lamina neurons (Ay-gal4 + 27G05-FLP) and LN are visualized with UAS-myr-RFP (magenta). (O) Quantification of the number of synapses in the distal region in *beat* single mutants (n > 102 axons). *beat-IIa or beat-IIb* single mutant did not show ectopic synapse formation in the distal region, indicating they work redundantly. (P) Quantification of the number of synapses in the distal region of lamina neurons upon simultaneous loss of *beat-IIa* and *beat-IIb* (n > 129 axons). Simultaneous downregulation of *beat-IIa* and *beat-IIb* showed ectopic synapse formation in the distal region. Statistical processing of the number of the number of synapses in the distal region of the lamina neuron was analyzed with the Mann-Whitney test.

**Figure S7.**
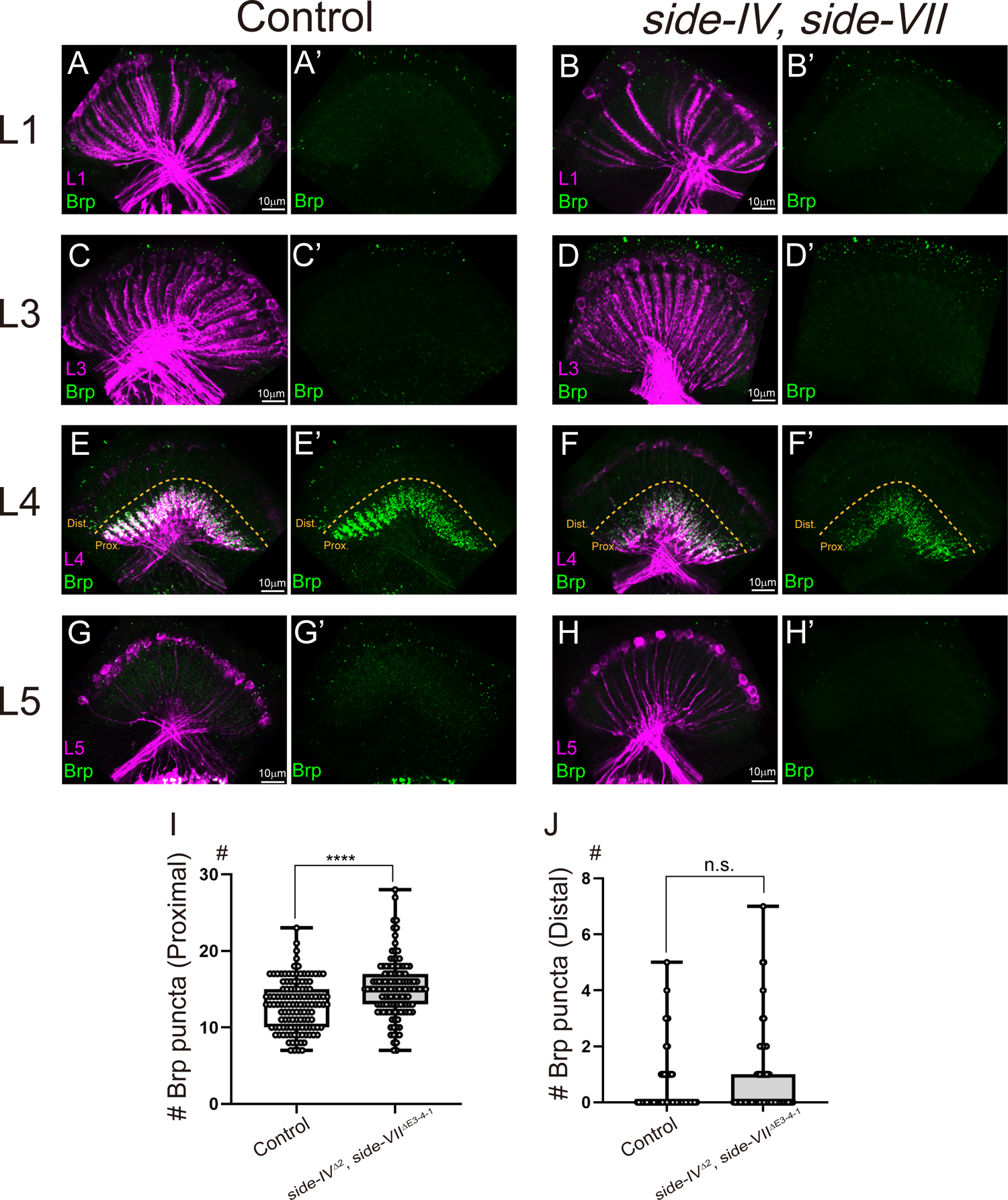
*side-IV* and *side-VII* double mutants do not form ectopic synapses in the distal region of L1, L3, L4, and L5 neurons (related to Figure 5) (A–H’) Axonal and synaptic morphology of each sub-type lamina neuron. Brp puncta (green) are visualized with STaR technique by cell-specific FLPase expression in each sub-type of lamina neurons. Morphology of lamina neurons are visualized with LexAop-myrtdTOM (magenta). (A–B’) L1 neuron. Control (A and A’), *side-IV^Δ2^*/*side-VII^ΔE3-4-1^* double mutant (B and B’) expressing FLPase specifically in the L1 neuron (48A08-AD (II), 66A01-DBD (III) L1 split-gal4 + UAS-FLP). (C–D’) L3 neuron. Control (C and C’), *side-IV^Δ2^*/*side-VII^ΔE3-4-1^*double mutant (D and D’) expressing FLPase specifically in L3 neuron (64B03-AD (II), 14B07-DBD (III) L3 split-gal4 + UAS-FLP). (E–F’) L4 neuron. Control (E and E’), *side-IV^Δ2^*/*side-VII^ΔE3-4-1^* double mutant (F and F’) expressing FLPase specifically in L4 neuron (31C06-AD (II), 34G07-DBD (III) L4 split-gal4 + UAS-FLP). Orange dotted lines indicate the boundary between the proximal and distal region. (G–H’) L5 neuron. Control (G and G’), *side-IV^Δ2^*/*side-VII^ΔE3-4-1^*double mutant (H and H’) expressing FLPase specifically in L5 neuron (64D07-AD (II), 37E10-DBD (III) L5 split-gal4 + UAS-FLP). (I) Quantification of the number of synapses in the proximal region of the L4 neuron (n > 123 axons). (J) Quantification of the number of synapses in the distal region of the L4 neuron (n > 191 axons). Statistical treatment of the number of the number of synapses in the proximal region and distal region of the L4 neuron were analyzed with the unpaired t-test and Mann-Whitney test, respectively.

**Figure S8.**
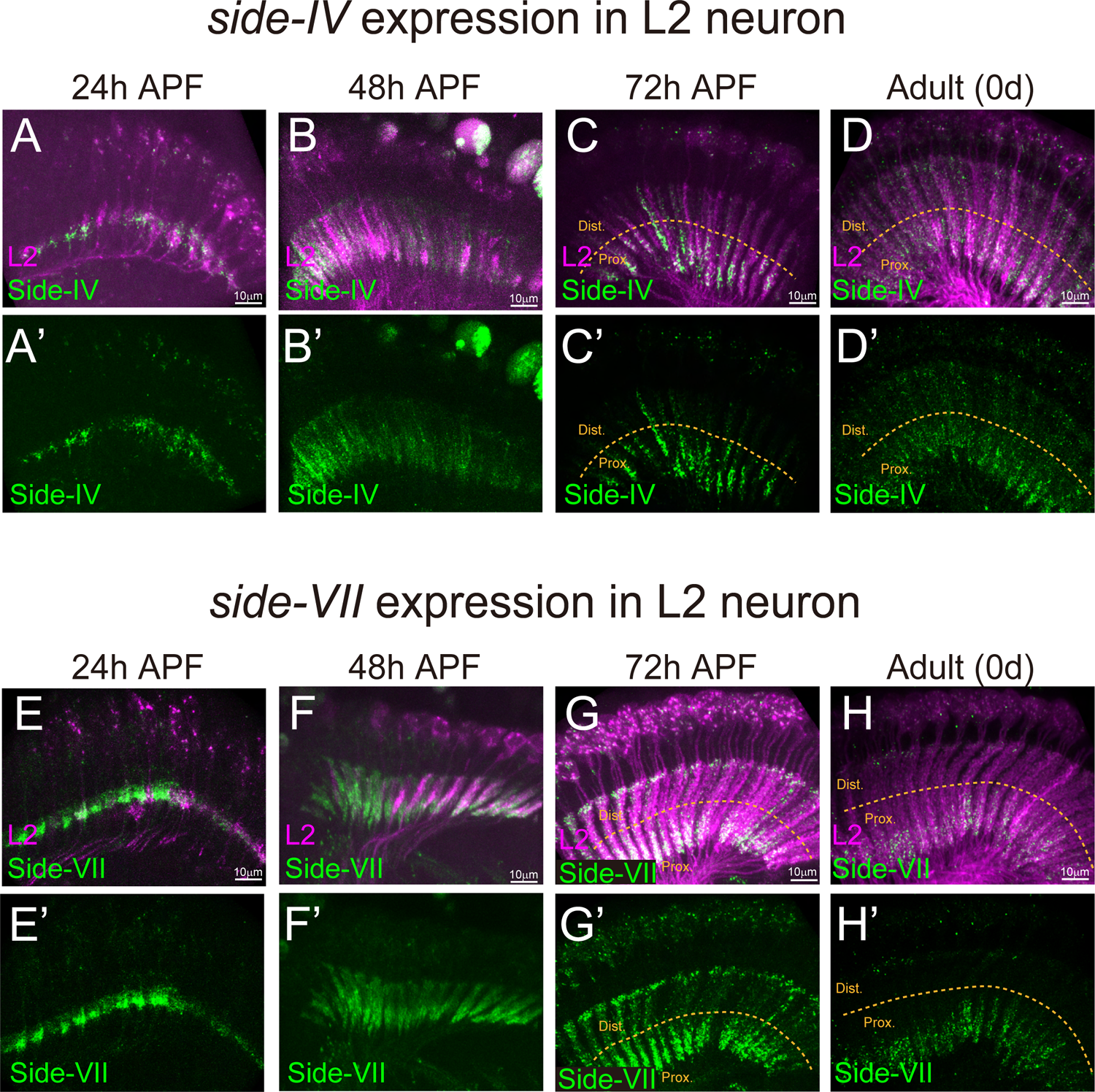
L2 neurons express *side-IV* and *side-VII* from developmental to adult (related to Figure 6) (A–H’) Endogenous expression of *side-IV* or *side-VII* in L2 neuron. Side-IV and Side-VII (green) are visualized by combining UAS-FLP with L2-gal4 (GMR16H03-gal4) or L2-split-gal4 (53G02-AD (II), 29G11-DBD (III)) against *side-IV*-FsF-GFP or *side-VII*-FsF-GFP. Morphology of L2 neurons are visualized with UAS-myr-RFP (magenta). *side-IV* expression in L2 neuron at 24 h APF (A and A’), 48 h APF (B and B’), 72 h APF (C and C’), 0d after eclosion (D and D’). Dotted lines indicate the boundary between the proximal and distal region. *side-VII* expression in L2 neuron at 24 h APF (E and E’), 48 h APF (F and F’), 72 h APF (G and G’), 0d after eclosion (H and H’).

## SUPPLEMENTALY TABLE

**Table S1**, The list of oligonucleotides used in this study

**Table S2**, The list of genotypes of flies used in this study

**Table S3**, The list of mutant alleles generated in this study

## RESOURCE AVAILABILITY

### Lead Contact

Further information and requests for resources and reagents should be directed to and will be fulfilled by the Lead Contact, Takashi Suzuki.

### Material Availability

All unique reagents generated in this study are available from the Lead Contact without restriction.

## METHOD DETAILS

### Key Resource Table

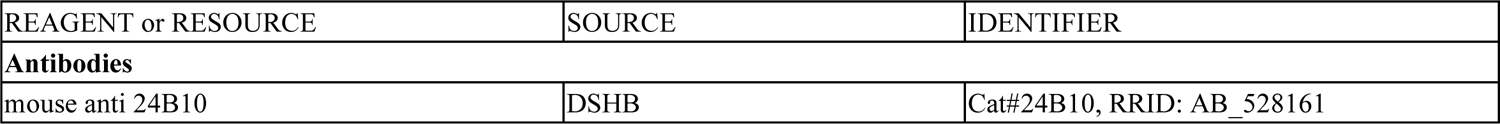
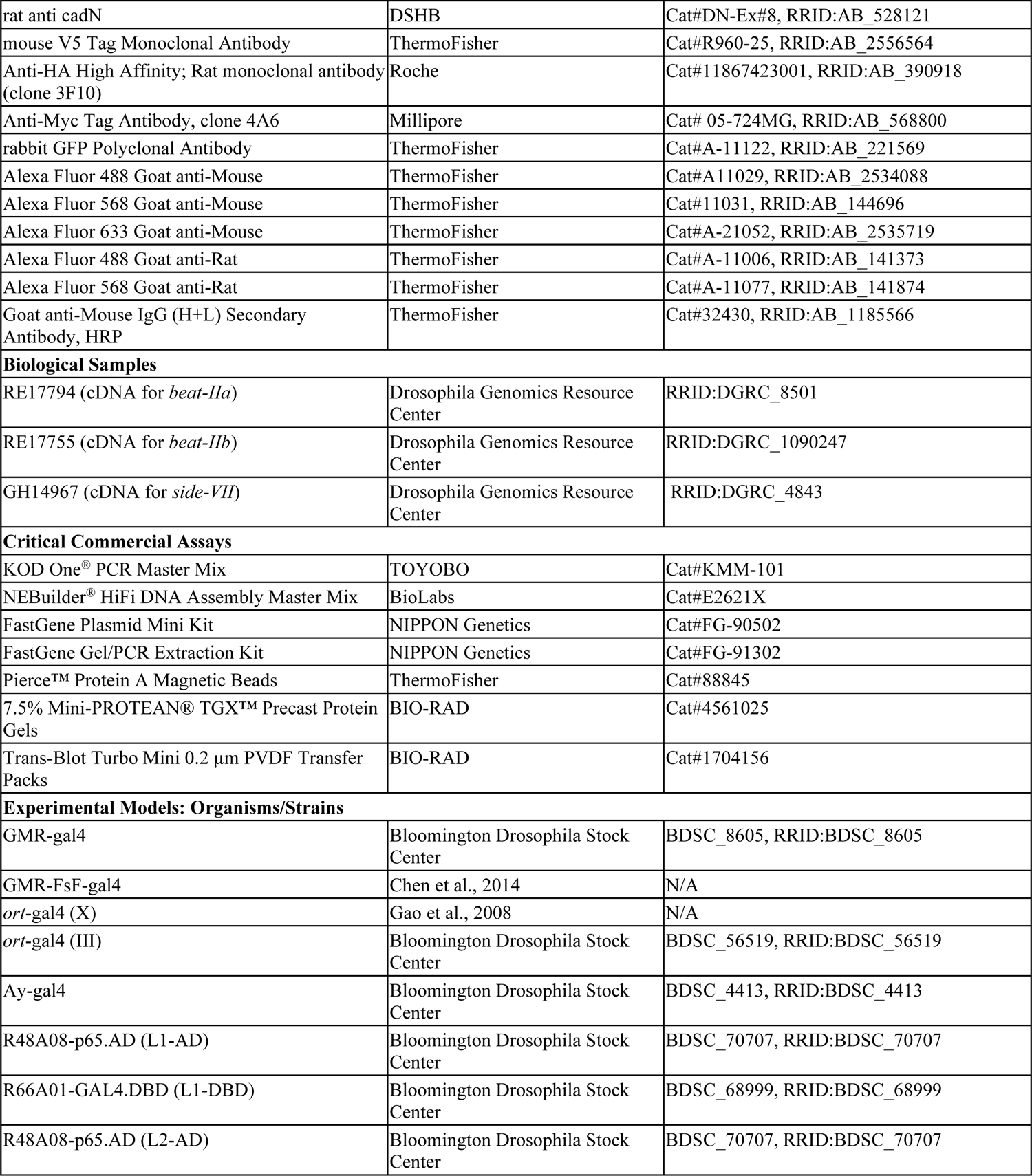
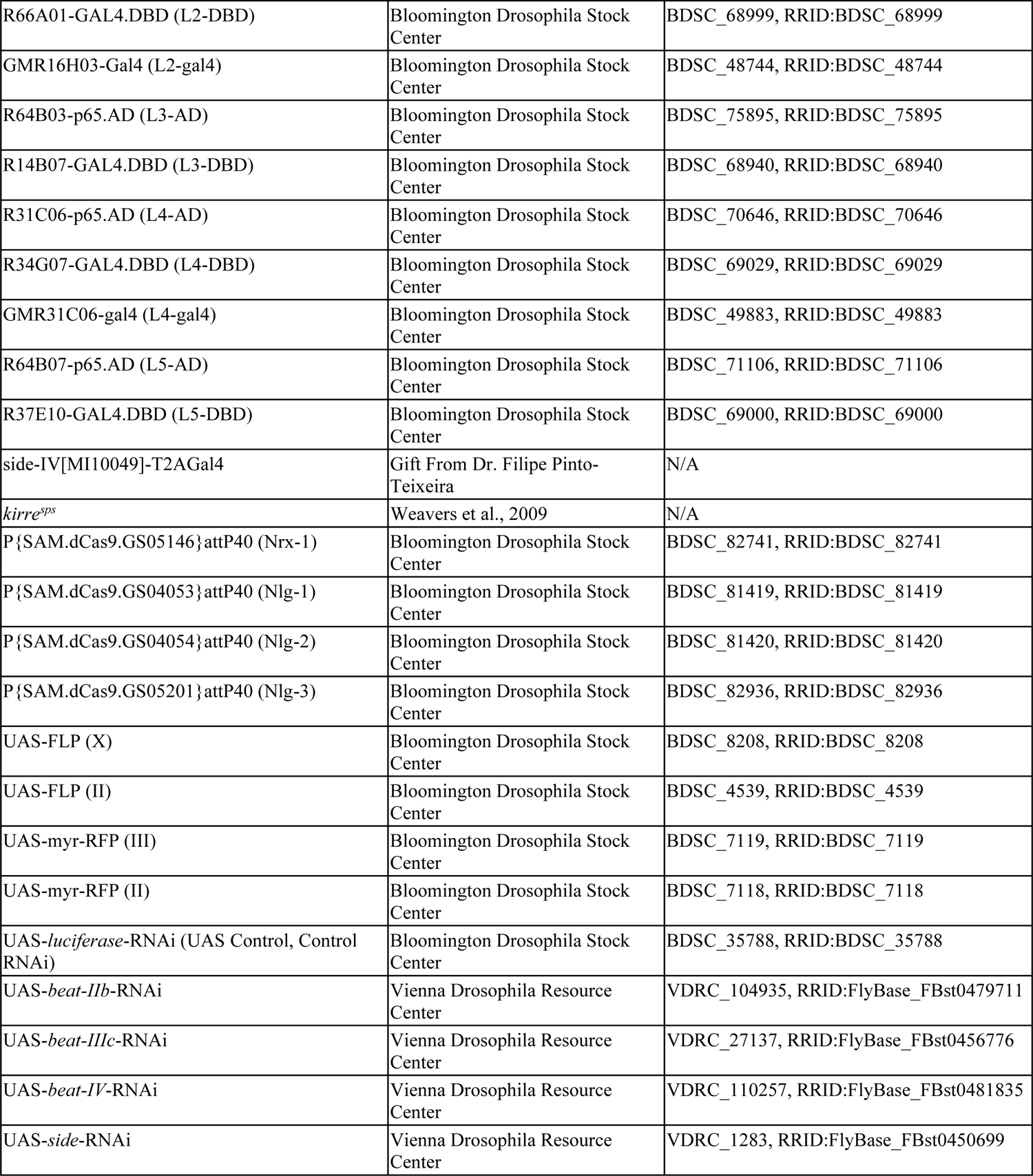
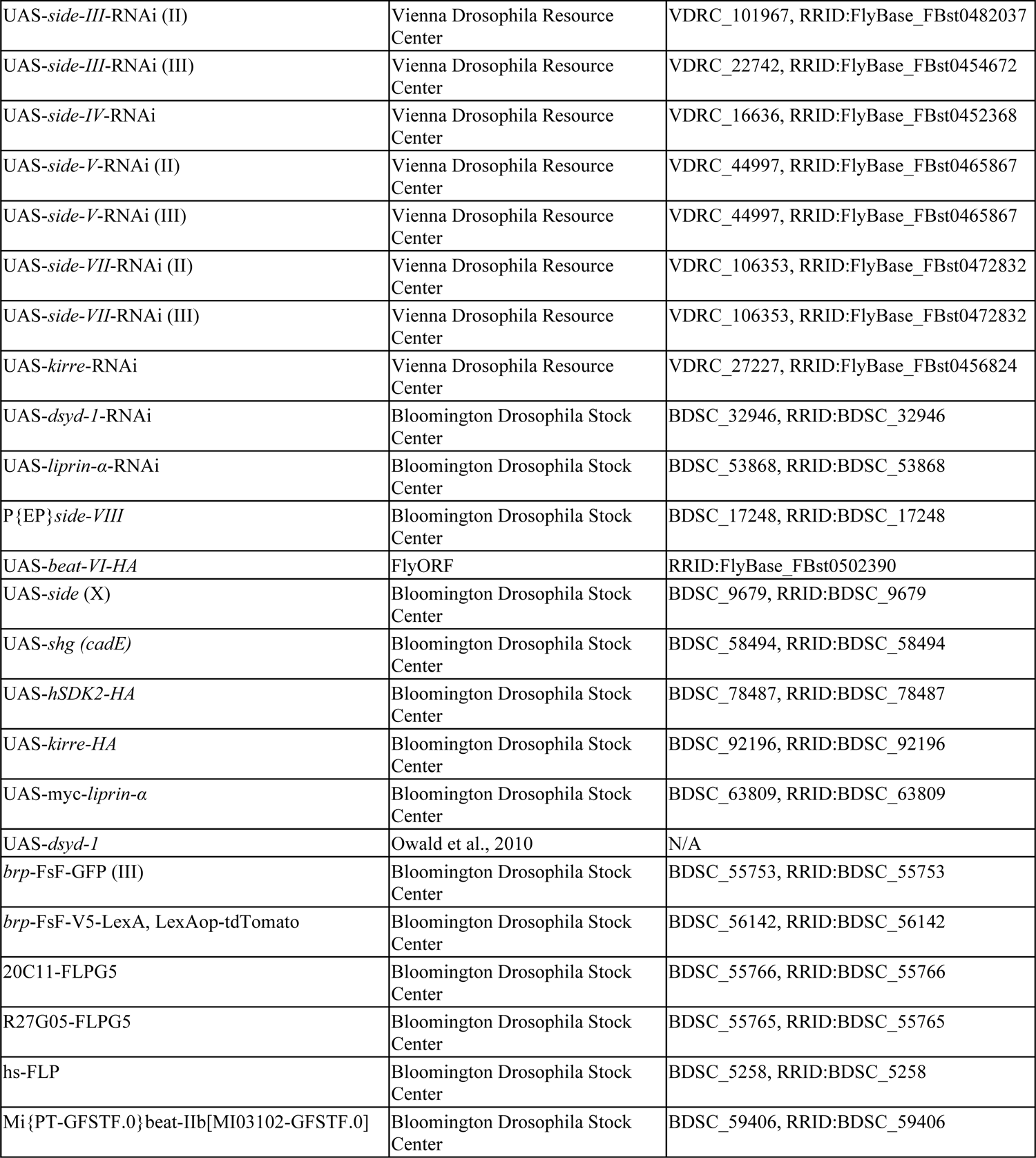
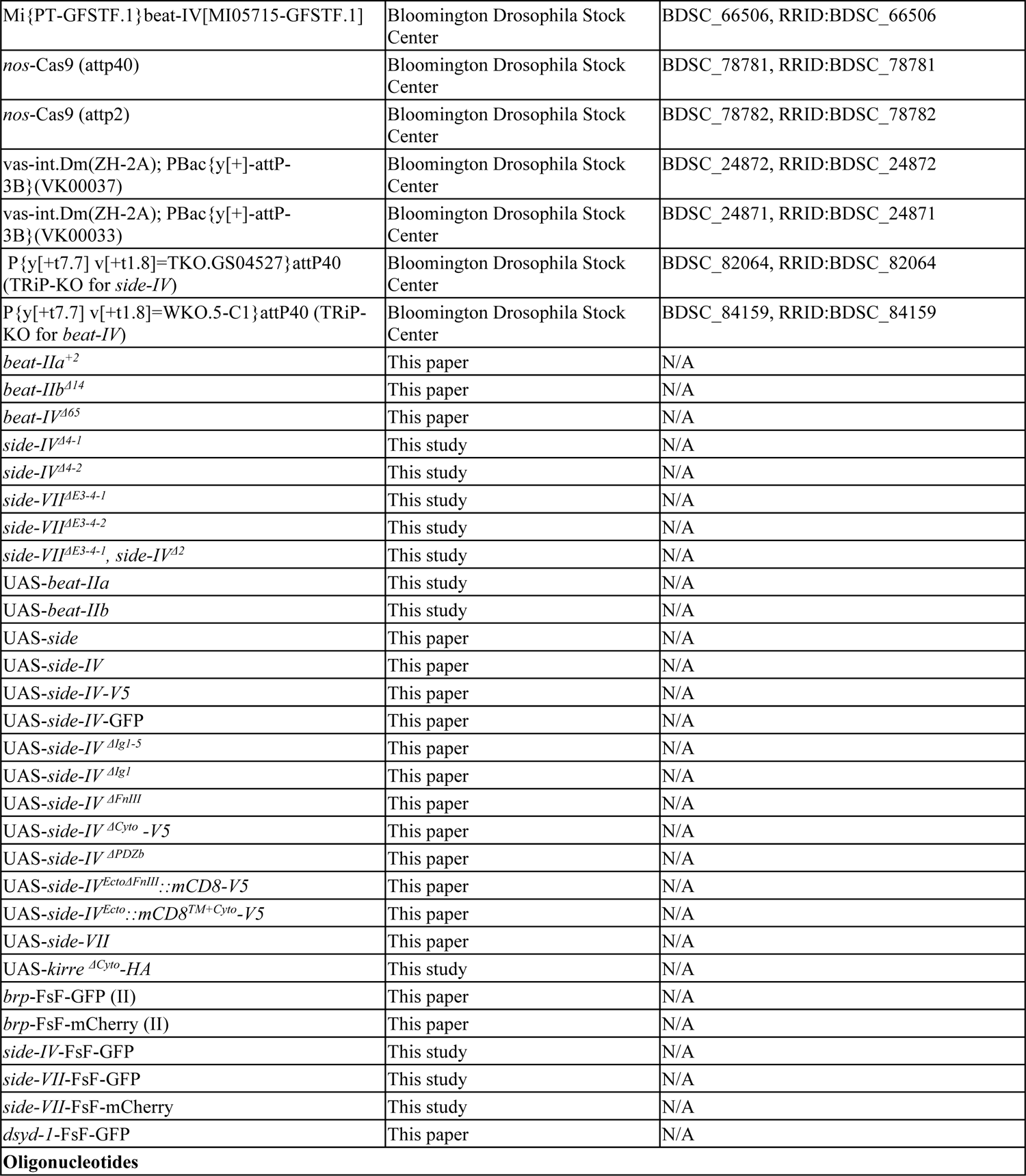
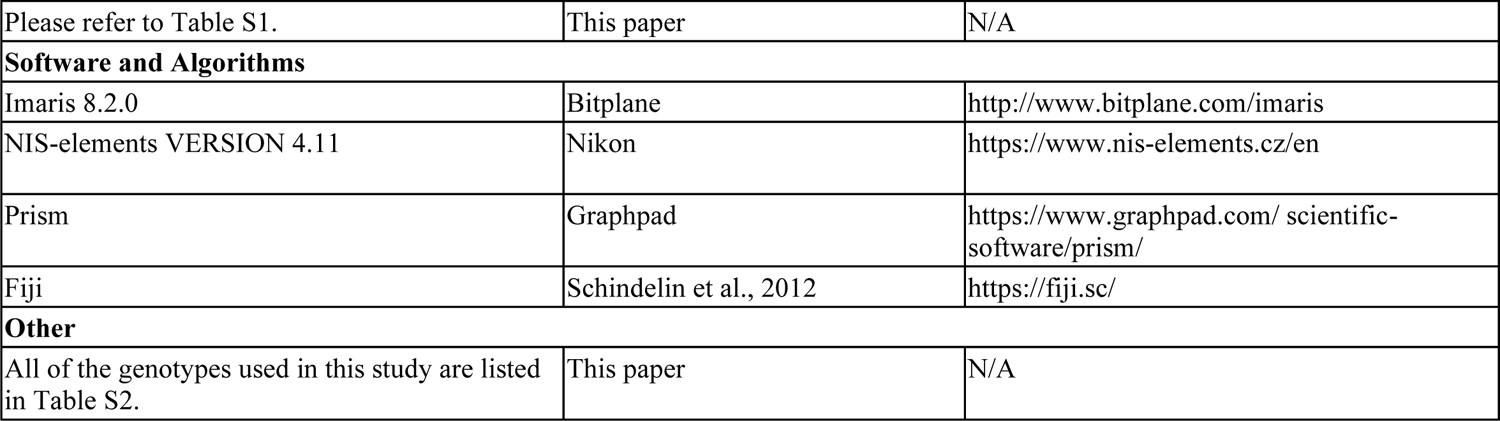

### Generation of transgenic flies

*side-IV*-FsF-GFP, *side-VII*-FsF-GFP (mCherry), *brp*-FsF-GFP (mCherry), and *dsyd-1*-FsF-GFP knock-in allele were generated by CRISPR/Cas9 technology.57 A knock-in vector containing the homology arms, the flip-out cassette (FRT-stop-FRT-GFP or mCherry), were generated as described previously.58 The oligo DNAs used for amplification of *side-IV*, *side-VII*, *brp*, and *dsyd-1* fragments and creating gDNA are listed in **Table S1**. A gRNA vectors were created and cloned into pBFv-U6.2 vector and were injected to eggs of nos-Cas9 (attp40) or nos-Cas9 (attp2) flies at WellGenetics or Bestgene together with the knock-in vector. The precise integration of the knock-in vector was verified by PCR and sequencing.

Mutant alleles of *beat-IIa*, *beat-IIb*, *beat-IV*, *side-IV*, and *side-VII* were generated by CRISPR/Cas9 technology.57 The gRNA vectors were created and cloned into pBFv-U6.2 vector for *beat-IIa*, *beat-IIb*, and *side-VII*. TRiP-KO lines (BDSC 84159, BDSC 82064) were used to generate mutant alleles of *beat-IV* and *side-IV*. For generating *side-IV*/*side-VII* double mutant, we introduced frameshift mutation *side-IV^2^* on *side-VII^E3-4-1^* allele by using TRiP-KO line (BDSC 82064). The oligo DNAs used for generating mutant alleles and all the mutations of mutant alleles are shown in **Table S1** and **Table S3**, respectively.

UAS lines were generated φC31 integrase-mediated recombination.^59^ The cDNAs were cloned into pUAST-attB vector and injected to egg of vas-int.Dm(ZH-2A); PBac{y[+]-attP-3B}(VK00037) or vasint. Dm(ZH-2A); PBac{y[+]-attP-3B}(VK00033) at Wellgenetics. The full length of cDNAs without UTRs were amplified from cDNA clones (RE17794 for *beat-IIa*, RE17755 for *beat-IIb*, ECD01053 for *side-IV,* GH14967 for *side-VII*) or a fly (BDSC 9679 for *side*). The available cDNA clones of *beat-IIb* (RE17755) and *side-IV* (ECD01053) did not cover all of the coding sequences. We amplified the lacking sequences from the fly genome and conjected the full length of cDNAs. We conjected 2 fragments (F1 & F2) by overlap pcr for generating insertions of Side-IV domain deletion variants or Side-IV-mCD8 chimeras. We referred to the beginning and end of the amino acid sequence of each domain of Side-IV, as previously reported.29 To generate the V5 or GFP tagged Side-IV constructs, GS-linker were inserted before the V5 or GFP. Furthermore, the sequence of PDZb motif of Side-IV was inserted in after the V5 or GFP not to disrupt the function of PDZb motif. The primers used to generate the UAS lines are shown in **Table S1**.

### Immunohistochemistry

The flies were maintained under standard conditions in Drosophila media at a temperature of 25°C. Experimental procedures for brain dissection, fixation, and immunostaining were performed as described previously.^60^ Fly brains were dissected in 0.1% PBT (PBS containing Triton X-100) and then fixed with 4% paraformaldehyde at room temperature for 60 min. The samples were washed with 0.1 % PBT 3 times and incubated at room temperature for 60 min. A primary antibody was added and incubated overnight at 4 degrees. The next day, thes samples were washed with 0.1 % PBT 3 times and incubated at room temperature for 60 min. The secondary antibody was added and incubated at room temperature for 60 min. Samples were washed with 0.1% PBT 3 times and incubated at room temperature for 2 hr. Finally, samples were washed with PBS once and replaced by VECTASHIELD®. For S2 cells, 1×10^5^ of cells were allowed to attach to glass slide at room temperature for 60 min before the fixation. The following antibodies were used for immunohistochemistry: mAb24B10 (1:50, DSHB), rat antibody to CadN (Ex#8, 1:50, DSHB), mouse antibody to V5 (R960, 1:400, Invitrogen), rat antibody to HA (3F10, 1:50, Roche), rat antibody to Myc (4A6, 1:50, Millipore). To enhance the signal intensity of Beat-IIb-MiMIC-GFP and Beat-IV-MiMIC-GFP, rabbit antibody to GFP conjugated with Alexa488 (1:400, Life technologies) was used. The secondary antibodies were Alexa488-, Alexa568- or Alexa633-conjugated (1:400, Life Technologies).

### Image acquisition

All confocal images were taken by Nikon C2+ or A1 confocal microscope with NIS elements (version 4.11). Each brain image was acquired between the depths of 30 and 50 μm from the surface of brain with a Z step size of 0.5 or 1μm. Images were acquired twice and averaged. Images were processed with NIS elements (Nikon), Adobe Photoshop, and Adobe illustrator.

### S2 cell transfection, aggregation assay, and co-immunoprecipitation

We used a mixture of culture medium; 500mL of Schneider’s Drosophila Medium (×1, gibco), 5mL of Penicillin-Streptomycin Solution (×100, wako), and 50mL of fetal bovine serum (Sigma). S2 cells were grown in 24 well plates (TrueLine) with 500μL of the culture medium. For transfection, 1.5μg of UAS vectors and pActin5C-gal4 were co-transfected with 10μL of HilyMax (DOJINDO) and 30μL of Schneider’s Drosophila Medium and incubated. For cell aggregation assay, S2 cells 48h after transfection were diluted to concentration of 105 cells ml-1 and agitated on a rotary shaker for 2 hours before the fixation. For co-immunoprecipitation, 7.5×105 of S2 cells were lysed in Lysis Buffer (50 mM Tris at pH 6.8, 150 mM NaCl, 1% NP40, protease inhibitor cocktail [Roche]), immunoprecipitated with 1 μg of GFP antibody (Life Technologies) and Protein A beads (Pierce) according to the manufacture’s procedure. SDS-PAGE was performed on 7.5% SDS-PAGE gels (Bio-Rad). Immunoblotting was performed by V5 antibody (R960, 1:2500, Invitrogen) and HRP secondary antibody (REF32430, 1:1000, Invitrogen). The images were taken by ImageQuant LAS 4000mini (cytiva) with Chemiluminescent HRP Substrate (MILLIPORE).

UAS plasmids used in S2 cells were generated by cloning cDNAs into pUAST-GFP or mCherry vectors. To amplify cDNA or domain deletion variants of *side-IV*, we used the same primers as generate transgenic flies. Genome extractions were performed to amplify the cDNA of *kirre* (UAS*-kirre.C-HA,* BDSC 92196), *cadE* (UASp-*shg.R*, BDSC58494), *dsyd-1* (UAS-*dsyd-1*),^61^ and *liprin-*μ (UAS-MYC-*liprin-*μ*.H*, BDSC63809). All of primers used to generate the UAS plasmids are shown in **Table S1**.

### QUANTIFICATION AND STATISTICAL ANALYSIS

All of the names of samples were randomized before the quantification. The number of Brp puncta of lamina neurons and photoreceptor R7 was counted manually with IMARIS (version 8.2.0) as described previously.^62^ The Brp puncta on a 3D image were identified with the spot function of IMARIS (diameter was set to 0.35μm). For the separation of the region of proximal region and distal region of lamina neurons, we cropped each stack of the proximal and distal ROI with NIS elements and a pen tablet (Cintiq 13HD, Wacom). We first drew the lines on the top and bottom of the lamina cartridge. Next, we drew a line on the midpoint of the lamina cartridge referring to the top and bottom lines. The accept region function of NIS-elements was then used to select the regions above and below the midpoint lines, which were then cropped as the proximal and distal regions, respectively. The number of Dsyd-1 puncta in the proximal region and distal region at 72 h APF were counted as the same method. The rate of overshoot axon was calculated manually with NIS elements, by counting the number of photoreceptors that overshoot beyond M6 layer and dividing the total number of axons.

In Figure 1M, the Side-IV cluster with Beat-IIb cluster was calculated manually with IMARIS, by counting the number of Side-IV puncta that co-localized with Beat-IIb puncta and dividing the total number of Side-IV puncta in the M1–M3 layer. The Side-IV/Beat-IIb cluster with Brp was calculated, by counting the number of Side-IV puncta co-localized with Beat-IIb and Brp puncta and dividing the total number of Side-IV puncta colocalized with Beat-IIb puncta.

In Figures 2H, I, and S2K the rate of Side-IV or Beat-IIb clustered axon were calculated manually with IMARIS, by the counting the number of axons with Side-IV or Beat-IIb clustered signal in the M1–M3 layer and dividing the total number of axons.

In Figures 3T and U, the cell-cell contact accumulation ratio was calculated manually with NIS-elements as described previously.43 Calculate the average intensity of GFP or mCherry signal on cell-cell border by drawing the line ROI and dividing the average intensity on the membrane, which did not contact other cells.

In Figures 4I and M, Pearson’s Coefficient was calculated with the JACoP plugin on Fiji.^63,64^

In Figures 6F and K, the rate of Side-IV or Side-VII positive axon was calculated manually with NIS elements, by counting the number of the axon that has Side-IV or Side-VII signal to the proximal region and dividing the total number of axons.

In Figure 7B, the average intensity along the Proximal to Distal regions of Side-IV and Kirre in the L2 neuron was calculated by drawing a linear ROI using NIS-elements and calculating the average intensity.

In Figure 7O, the rate of Dsyd-1 positive axon (Distal) was calculated manually with NIS-elements, by counting the number of the axon that has Dsyd-1 signal on the distal region and dividing the total number of the axon at 48 h APF.

In Figure S2Q, the number of cells per cell aggregation was calculated manually with NIS-elements, by counting the total number of cells on each cell aggregation. The cell cluster with contains both GFP and mCherry expressing cells was identified as cell aggregation.

In Figure S4C, the rate of Kirre-GFP cell cluster with and without mCherry positive cell was calculated manually with NIS-elelments, by counting the rate of Kirre-GFP cell cluster which had more than one mCherry positive cell. The cell cluster with more than two GFP expressing cells was identified as Kirre-GFP cell cluster.

## Statistical analysis

All of the statistical analysis are described in the figure legends. Statistical significance. n.s. *P* > 0.05, * *P* ≤ 0.05, ** *P* ≤ 0.01, *** *P* ≤ 0.001, **** *P* ≤ 0.0001.

## REFERENCES

1. Sperry, R.W. (1963). CHEMOAFFINITY IN THE ORDERLY GROWTH OF NERVE FIBER PATTERNS AND CONNECTIONS. Proceedings of the National Academy of Sciences of the United States of America 50, 703–710. 10.1073/pnas.50.4.703.

2. Sanes, J.R., and Zipursky, S.L. (2020). Synaptic Specificity, Recognition Molecules, and Assembly of Neural Circuits. Cell 181, 536–556. 10.1016/j.cell.2020.04.008.

3. Özel, M.N., Kulkarni, A., Hasan, A., Brummer, J., Moldenhauer, M., Daumann, I.M., Wolfenberg, H., Dercksen, V.J., Kiral, F.R., Weiser, M., et al. (2019). Serial Synapse Formation through Filopodial Competition for Synaptic Seeding Factors. Developmental cell 50, 447–461.e448. 10.1016/j.devcel.2019.06.014.

4. Kiral, F.R., Linneweber, G.A., Mathejczyk, T., Georgiev, S.V., Wernet, M.F., Hassan, B.A., von Kleist, M., and Hiesinger, P.R. (2020). Autophagy-dependent filopodial kinetics restrict synaptic partner choice during Drosophila brain wiring. Nature communications 11, 1325. 10.1038/s41467-020-14781-4.

5. Hiesinger, P.R. (2021). Brain wiring with composite instructions. BioEssays: news and reviews in molecular, cellular and developmental biology 43, e2000166. 10.1002/bies.202000166.

6. Südhof, T.C. (2021). The cell biology of synapse formation. The Journal of cell biology 220. 10.1083/jcb.202103052.

7. Xu, C., Theisen, E., Maloney, R., Peng, J., Santiago, I., Yapp, C., Werkhoven, Z., Rumbaut, E., Shum, B., Tarnogorska, D., et al. (2019). Control of Synaptic Specificity by Establishing a Relative Preference for Synaptic Partners. Neuron 103, 865–877.e867. 10.1016/j.neuron.2019.06.006.

8. Kiral, F.R., Dutta, S.B., Linneweber, G.A., Hilgert, S., Poppa, C., Duch, C., von Kleist, M., Hassan, B.A., and Hiesinger, P.R. (2021). Brain connectivity inversely scales with developmental temperature in Drosophila. Cell Reports 37, 110145. https://doi.org/10.1016/j.celrep.2021.110145.

9. Takemura, S.Y., Bharioke, A., Lu, Z., Nern, A., Vitaladevuni, S., Rivlin, P.K., Katz, W.T., Olbris, D.J., Plaza, S.M., Winston, P., et al. (2013). A visual motion detection circuit suggested by Drosophila connectomics. Nature 500, 175–181. 10.1038/nature12450.

10. Takemura, S.-y., Xu, C.S., Lu, Z., Rivlin, P.K., Parag, T., Olbris, D.J., Plaza, S., Zhao, T., Katz, W.T., Umayam, L., et al. (2015). Synaptic circuits and their variations within different columns in the visual system of Drosophila. Proceedings of the National Academy of Sciences of the United States of America 112, 13711–13716. 10.1073/pnas.1509820112.

11. Davis, F.P., Nern, A., Picard, S., Reiser, M.B., Rubin, G.M., Eddy, S.R., and Henry, G.L. (2020). A genetic, genomic, and computational resource for exploring neural circuit function. eLife 9, e50901. 10.7554/eLife.50901.

12. Cosmanescu, F., Katsamba, P.S., Sergeeva, A.P., Ahlsen, G., Patel, S.D., Brewer, J.J., Tan, L., Xu, S., Xiao, Q., Nagarkar-Jaiswal, S., et al. (2018). Neuron-Subtype-Specific Expression, Interaction Affinities, and Specificity Determinants of DIP/Dpr Cell Recognition Proteins. Neuron 100, 1385–1400.e1386. 10.1016/j.neuron.2018.10.046.

13. Kurmangaliyev, Y.Z., Yoo, J., Valdes-Aleman, J., Sanfilippo, P., and Zipursky, S.L. (2020). Transcriptional Programs of Circuit Assembly in the Drosophila Visual System. Neuron 108, 1045–1057.e1046. 10.1016/j.neuron.2020.10.006.

14. Özel, M.N., Simon, F., Jafari, S., Holguera, I., Chen, Y.C., Benhra, N., El-Danaf, R.N., Kapuralin, K., Malin, J.A., Konstantinides, N., and Desplan, C. (2021). Neuronal diversity and convergence in a visual system developmental atlas. Nature 589, 88–95. 10.1038/s41586-020-2879-3.

15. Jain, S., Lin, Y., Kurmangaliyev, Y.Z., Valdes-Aleman, J., LoCascio, S.A., Mirshahidi, P., Parrington, B., and Zipursky, S.L. (2022). A global timing mechanism regulates cell-type-specific wiring programmes. Nature 603, 112–118. 10.1038/s41586-022-04418-5.

16. Özkan, E., Carrillo, R.A., Eastman, C.L., Weiszmann, R., Waghray, D., Johnson, K.G., Zinn, K., Celniker, S.E., and Garcia, K.C. (2013). An extracellular interactome of immunoglobulin and LRR proteins reveals receptor-ligand networks. Cell 154, 228–239. 10.1016/j.cell.2013.06.006.

17. Tan, L., Zhang, K.X., Pecot, M.Y., Nagarkar-Jaiswal, S., Lee, P.-T., Takemura, S.-Y., McEwen, J.M., Nern, A., Xu, S., Tadros, W., et al. (2015). Ig Superfamily Ligand and Receptor Pairs Expressed in Synaptic Partners in Drosophila. Cell 163, 1756–1769. 10.1016/j.cell.2015.11.021.

18. Carrillo, R.A., Özkan, E., Menon, K.P., Nagarkar-Jaiswal, S., Lee, P.T., Jeon, M., Birnbaum, M.E., Bellen, H.J., Garcia, K.C., and Zinn, K. (2015). Control of Synaptic Connectivity by a Network of Drosophila IgSF Cell Surface Proteins. Cell 163, 1770–1782. 10.1016/j.cell.2015.11.022.

19. Sergeeva, A.P., Katsamba, P.S., Cosmanescu, F., Brewer, J.J., Ahlsen, G., Mannepalli, S., Shapiro, L., and Honig, B. (2020). DIP/Dpr interactions and the evolutionary design of specificity in protein families. Nature communications 11, 2125. 10.1038/s41467-020-15981-8.

20. Barish, S., Nuss, S., Strunilin, I., Bao, S., Mukherjee, S., Jones, C.D., and Volkan, P.C. (2018). Combinations of DIPs and Dprs control organization of olfactory receptor neuron terminals in Drosophila. PLoS genetics 14, e1007560. 10.1371/journal.pgen.1007560.

21. suppXu, S., Xiao, Q., Cosmanescu, F., Sergeeva, A.P., Yoo, J., Lin, Y., Katsamba, P.S., Ahlsen, G., Kaufman, J., Linaval, N.T., et al. (2018). Interactions between the Ig-Superfamily Proteins DIP-α and Dpr6/10 Regulate Assembly of Neural Circuits. Neuron 100, 1369–1384.e1366. 10.1016/j.neuron.2018.11.001.

22. Courgeon, M., and Desplan, C. (2019). Coordination between stochastic and deterministic specification in the Drosophila visual system. Science (New York, N.Y.) 366. 10.1126/science.aay6727.

23. Venkatasubramanian, L., Guo, Z., Xu, S., Tan, L., Xiao, Q., Nagarkar-Jaiswal, S., and Mann, R.S. (2019). Stereotyped terminal axon branching of leg motor neurons mediated by IgSF proteins DIP-α and Dpr10. eLife 8, e42692. 10.7554/eLife.42692.

24. Bornstein, B., Meltzer, H., Adler, R., Alyagor, I., Berkun, V., Cummings, G., Reh, F., Keren-Shaul, H., David, E., Riemensperger, T., and Schuldiner, O. (2021). Transneuronal Dpr12/DIP-δ interactions facilitate compartmentalized dopaminergic innervation of Drosophila mushroom body axons. The EMBO journal 40, e105763. 10.15252/embj.2020105763.

25. Vactor, D.V., Sink, H., Fambrough, D., Tsoo, R., and Goodman, C.S. (1993). Genes that control neuromuscular specificity in Drosophila. Cell 73, 1137–1153. 10.1016/0092-8674(93)90643-5.

26. Fambrough, D., and Goodman, C.S. (1996). The Drosophila beaten path gene encodes a novel secreted protein that regulates defasciculation at motor axon choice points. Cell 87, 1049–1058. 10.1016/s0092-8674(00)81799-7.

27. Sink, H., Rehm, E.J., Richstone, L., Bulls, Y.M., and Goodman, C.S. (2001). sidestep encodes a target-derived attractant essential for motor axon guidance in Drosophila. Cell 105, 57–67. 10.1016/s0092-8674(01)00296-3.

28. Siebert, M., Banovic, D., Goellner, B., and Aberle, H. (2009). Drosophila motor axons recognize and follow a Sidestep-labeled substrate pathway to reach their target fields. Genes Dev 23, 1052–1062. 10.1101/gad.520509.

29. Li, H., Watson, A., Olechwier, A., Anaya, M., Sorooshyari, S.K., Harnett, D.P., Lee, H.-K.P., Vielmetter, J., Fares, M.A., Garcia, K.C., et al. (2017). Deconstruction of the beaten Path-Sidestep interaction network provides insights into neuromuscular system development. eLife 6, e28111. 10.7554/eLife.28111.

30. Pipes, G.C., Lin, Q., Riley, S.E., and Goodman, C.S. (2001). The Beat generation: a multigene family encoding IgSF proteins related to the Beat axon guidance molecule in Drosophila. Development (Cambridge, England) 128, 4545–4552. 10.1242/dev.128.22.4545.

31. Aberle, H. (2009). Searching for guidance cues: follow the Sidestep trail. Fly 3, 270–273. 10.4161/fly.9790.

32. Zinn, K. (2009). Choosing the road less traveled by: a ligand-receptor system that controls target recognition by Drosophila motor axons. Genes & development 23, 1042–1045. 10.1101/gad.1803009.

33. Kinold, J.C., Pfarr, C., and Aberle, H. (2018). Sidestep-induced neuromuscular miswiring causes severe locomotion defects in Drosophila larvae. Development (Cambridge, England) 145. 10.1242/dev.163279.

34. Kind, E., Longden, K.D., Nern, A., Zhao, A., Sancer, G., Flynn, M.A., Laughland, C.W., Gezahegn, B., Ludwig, H.D.F., Thomson, A.G., et al. (2021). Synaptic targets of photoreceptors specialized to detect color and skylight polarization in Drosophila. eLife 10, e71858. 10.7554/eLife.71858.

35. Heymann, C., Paul, C., Huang, N., Kinold, J.C., Dietrich, A.C., and Aberle, H. (2022). Molecular insights into the axon guidance molecules Sidestep and Beaten path. Frontiers in physiology 13, 1057413. 10.3389/fphys.2022.1057413.

36. Karuppudurai, T., Lin, T.-Y., Ting, C.-Y., Pursley, R., Melnattur, Krishna V., Diao, F., White, Benjamin H., Macpherson, Lindsey J., Gallio, M., Pohida, T., and Lee, C.-H. (2014). A Hard-Wired Glutamatergic Circuit Pools and Relays UV Signals to Mediate Spectral Preference in Drosophila. Neuron 81, 603–615. https://doi.org/10.1016/j.neuron.2013.12.010.

37. Chen, Y., Akin, O., Nern, A., Tsui, C.Y., Pecot, M.Y., and Zipursky, S.L. (2014). Cell-type-specific labeling of synapses in vivo through synaptic tagging with recombination. Neuron 81, 280–293. 10.1016/j.neuron.2013.12.021.

38. Nagarkar-Jaiswal, S., Lee, P.-T., Campbell, M.E., Chen, K., Anguiano-Zarate, S., Gutierrez, M.C., Busby, T., Lin, W.-W., He, Y., Schulze, K.L., et al. (2015). A library of MiMICs allows tagging of genes and reversible, spatial and temporal knockdown of proteins in Drosophila. eLife 4, e05338. 10.7554/eLife.05338.

39. Tanaka, H., Miyazaki, N., Matoba, K., Nogi, T., Iwasaki, K., and Takagi, J. (2012). Higher-order architecture of cell adhesion mediated by polymorphic synaptic adhesion molecules neurexin and neuroligin. Cell reports 2, 101–110. 10.1016/j.celrep.2012.06.009.

40. Gao, S., Takemura, S.Y., Ting, C.Y., Huang, S., Lu, Z., Luan, H., Rister, J., Thum, A.S., Yang, M., Hong, S.T., et al. (2008). The neural substrate of spectral preference in Drosophila. Neuron 60, 328–342. 10.1016/j.neuron.2008.08.010.

41. Fischbach, K.F., Linneweber, G.A., Andlauer, T.F., Hertenstein, A., Bonengel, B., and Chaudhary, K. (2009). The irre cell recognition module (IRM) proteins. Journal of neurogenetics 23, 48–67. 10.1080/01677060802471668.

42. Strutt, H., and Strutt, D. (2008). Differential stability of flamingo protein complexes underlies the establishment of planar polarity. Current biology: CB 18, 1555–1564. 10.1016/j.cub.2008.08.063.

43. Hakeda-Suzuki, S., Berger-Müller, S., Tomasi, T., Usui, T., Horiuchi, S.Y., Uemura, T., and Suzuki, T. (2011). Golden Goal collaborates with Flamingo in conferring synaptic-layer specificity in the visual system. Nature neuroscience 14, 314–323. 10.1038/nn.2756.

44. Owald, D., Khorramshahi, O., Gupta, V.K., Banovic, D., Depner, H., Fouquet, W., Wichmann, C., Mertel, S., Eimer, S., Reynolds, E., et al. (2012). Cooperation of Syd-1 with Neurexin synchronizes pre-with postsynaptic assembly. Nature neuroscience 15, 1219–1226. 10.1038/nn.3183.

45. Fouquet, W., Owald, D., Wichmann, C., Mertel, S., Depner, H., Dyba, M., Hallermann, S., Kittel, R.J., Eimer, S., and Sigrist, S.J. (2009). Maturation of active zone assembly by Drosophila Bruchpilot. The Journal of cell biology 186, 129–145. 10.1083/jcb.200812150.

46. Rivera-Alba, M., Vitaladevuni, S.N., Mishchenko, Y., Lu, Z., Takemura, S.Y., Scheffer, L., Meinertzhagen, I.A., Chklovskii, D.B., and de Polavieja, G.G. (2011). Wiring economy and volume exclusion determine neuronal placement in the Drosophila brain. Current biology: CB 21, 2000–2005. 10.1016/j.cub.2011.10.022.

47. Varoqueaux, F., Aramuni, G., Rawson, R.L., Mohrmann, R., Missler, M., Gottmann, K., Zhang, W., Südhof, T.C., and Brose, N. (2006). Neuroligins determine synapse maturation and function. Neuron 51, 741–754. 10.1016/j.neuron.2006.09.003.

48. Gokce, O., and Südhof, T.C. (2013). Membrane-tethered monomeric neurexin LNS-domain triggers synapse formation. The Journal of neuroscience: the official journal of the Society for Neuroscience 33, 14617–14628. 10.1523/jneurosci.1232-13.2013.

49. Fischbach, K.F., and Dittrich, A.P.M. (1989). The optic lobe of Drosophila melanogaster. I. A Golgi analysis of wild-type structure. Cell and Tissue Research 258, 441–475. 10.1007/BF00218858.

50. Kolodziejczyk, A., Sun, X., Meinertzhagen, I.A., and Nässel, D.R. (2008). Glutamate, GABA and acetylcholine signaling components in the lamina of the Drosophila visual system. PLoS One 3, e2110–e2110. 10.1371/journal.pone.0002110.

51. Lüthy, K., Ahrens, B., Rawal, S., Lu, Z., Tarnogorska, D., Meinertzhagen, I.A., and Fischbach, K.F. (2014). The irre cell recognition module (IRM) protein Kirre is required to form the reciprocal synaptic network of L4 neurons in the Drosophila lamina. Journal of neurogenetics 28, 291–301. 10.3109/01677063.2014.883390.

52. Weavers, H., Prieto-Sánchez, S., Grawe, F., Garcia-López, A., Artero, R., Wilsch-Bräuninger, M., Ruiz-Gómez, M., Skaer, H., and Denholm, B. (2009). The insect nephrocyte is a podocyte-like cell with a filtration slit diaphragm. Nature 457, 322–326. 10.1038/nature07526.

53. Kaufmann, N., DeProto, J., Ranjan, R., Wan, H., and Van Vactor, D. (2002). Drosophila liprin-alpha and the receptor phosphatase Dlar control synapse morphogenesis. Neuron 34, 27–38. 10.1016/s0896-6273(02)00643-8.

54. Jiang, X., Sando, R., and Südhof, T.C. (2021). Multiple signaling pathways are essential for synapse formation induced by synaptic adhesion molecules. Proceedings of the National Academy of Sciences of the United States of America 118. 10.1073/pnas.2000173118.

55. Lobb-Rabe, M., Nawrocka, W.I., Carrillo, R.A., and Özkan, E. (2023). Neuronal wiring receptors Dprs and DIPs are GPI anchored and this modification contributes to their cell surface organization. bioRxiv, 2023.2003.2002.530872. 10.1101/2023.03.02.530872.

56. Simon, F., and Konstantinides, N. (2021). Single-cell transcriptomics in the Drosophila visual system: Advances and perspectives on cell identity regulation, connectivity, and neuronal diversity evolution. Developmental biology 479, 107–122. 10.1016/j.ydbio.2021.08.001.

57. Kondo, S., and Ueda, R. (2013). Highly improved gene targeting by germline-specific Cas9 expression in Drosophila. Genetics 195, 715–721. 10.1534/genetics.113.156737.

58. Trush, O., Liu, C., Han, X., Nakai, Y., Takayama, R., Murakawa, H., Carrillo, J.A., Takechi, H., Hakeda-Suzuki, S., Suzuki, T., and Sato, M. (2019). N-Cadherin Orchestrates Self-Organization of Neurons within a Columnar Unit in the <em>Drosophila</em> Medulla. The Journal of Neuroscience 39, 5861–5880. 10.1523/jneurosci.3107-18.2019.

59. Thorpe, H.M., and Smith, M.C. (1998). In vitro site-specific integration of bacteriophage DNA catalyzed by a recombinase of the resolvase/invertase family. Proceedings of the National Academy of Sciences of the United States of America 95, 5505–5510. 10.1073/pnas.95.10.5505.

60. Hakeda-Suzuki, S., Takechi, H., Kawamura, H., and Suzuki, T. (2017). Two receptor tyrosine phosphatases dictate the depth of axonal stabilizing layer in the visual system. eLife 6, e31812. 10.7554/eLife.31812.

61. Owald, D., Fouquet, W., Schmidt, M., Wichmann, C., Mertel, S., Depner, H., Christiansen, F., Zube, C., Quentin, C., Körner, J., et al. (2010). A Syd-1 homologue regulates pre- and postsynaptic maturation in Drosophila. Journal of Cell Biology 188, 565–579. 10.1083/jcb.200908055.

62. Sugie, A., Hakeda-Suzuki, S., Suzuki, E., Silies, M., Shimozono, M., Möhl, C., Suzuki, T., and Tavosanis, G. (2015). Molecular Remodeling of the Presynaptic Active Zone of Drosophila Photoreceptors via Activity-Dependent Feedback. Neuron 86, 711–725. 10.1016/j.neuron.2015.03.046.

63. Bolte, S., and Cordelières, F.P. (2006). A guided tour into subcellular colocalization analysis in light microscopy. Journal of microscopy 224, 213–232. 10.1111/j.1365-2818.2006.01706.x.

64. Schindelin, J., Arganda-Carreras, I., Frise, E., Kaynig, V., Longair, M., Pietzsch, T., Preibisch, S., Rueden, C., Saalfeld, S., Schmid, B., et al. (2012). Fiji: an open-source platform for biological-image analysis. Nature methods 9, 676–682. 10.1038/nmeth.2019.

